# Evidence accumulation and change rate inference in dynamic environments

**DOI:** 10.1101/066480

**Authors:** Adrian E. Radillo, Alan Veliz-Cuba, Krešimir Josić, Zachary P. Kilpatrick

## Abstract

In a constantly changing world, animals must account for environmental volatility when making decisions. To appropriately discount older, irrelevant information, they need to learn the rate at which the environment changes. We develop an ideal observer model capable of inferring the present state of the environment along with its rate of change. Key to this computation is updating the posterior probability of all possible changepoint counts. This computation can be challenging, as the number of possibilities grows rapidly with time. However, we show how the computations can be simplified in the continuum limit by a moment closure approximation. The resulting low-dimensional system can be used to infer the environmental state and change rate with accuracy comparable to the ideal observer. The approximate computations can be performed by a neural network model via a rate-correlation based plasticity rule. We thus show how optimal observers accumulates evidence in changing environments, and map this computation to reduced models which perform inference using plausible neural mechanisms.

## 1 Introduction

Animals continuously make decisions in order to find food, identify mates, and avoid predators. However, the world is seldom static. Information that was critical yesterday may be of little value now. Thus, when accumulating evidence to decide on a course of action, animals weight new evidence more strongly than old (Pearson et al, 2011). The rate at which the world changes determines the rate at which an individual should discount previous information (Deneve, 2008; Veliz-Cuba et al, 2016). For instance, when actively tracking prey, a predator may only use visual information obtained within the last second (Olberg et al, 2000; Portugues and Engert, 2009), while social insect colonies integrate evidence that can be hours to days old when deciding on a new home site (Franks et al, 2002). However, the timescale at which the environment changes is unlikely to be known in advance. To make accurate decisions, animals must also learn how rapidly their environment changes (Wilson et al, 2010).

Evidence accumulators are often used to model decision processes in static and fluctuating environments (Smith and Ratcliff, 2004; Bogacz et al, 2006). These models show how noisy observations can be accumulated to provide a probability that one among multiple alternatives is correct (Gold and Shadlen, 2007; Beck et al, 2008). They explain a variety of behavioral data (Ratcliff and McKoon, 2008; Brunton et al, 2013), and electrophysiological recordings suggest that neural activity can reflect the accumulation of evidence (Huk and Shadlen, 2005; Kira et al, 2015). Since normative evidence accumulation models determine the belief of an ideal observer, they also show the best way to integrate noisy sensory measurements, and can tell us if and how animals fail to use such information optimally (Bogacz et al, 2006; Beck et al, 2008).

Early decision-making models focused on decisions between two choices in a static environment (Wald and Wolfowitz, 1948; Gold and Shadlen, 2007). Recent studies have extended this work to more ecologically relevant situations, including multiple alternatives (Churchland et al, 2008; Krajbich and Rangel, 2011), multidimensional environments (Niv et al, 2015), and cases where the correct choice (McGuire et al, 2014; Glaze et al, 2015), or context (Shvartsman et al, 2015), changes in time. In these cases, normative models are more difficult to derive and analyze (Wilson and Niv, 2011), and their dynamics are more complex. However, methods of sequential and stochastic analysis are still useful in understanding their properties (Wilson et al, 2010; Veliz-Cuba et al, 2016).

We examine the case of a changing environment where an optimal observer discounts prior evidence at a rate determined by environmental volatility. Experiments suggest that humans learn the rate of environmental fluctuations to make choices nearly optimally (Glaze et al, 2015). During dynamic foraging experiments where the choice with the highest reward changes in time, monkeys also appear to use an evidence discounting strategy suited to the environmental change rate (Sugrue et al, 2004).

However, most previous models have assumed that the rate of change of the environment is known ahead of time to the observer (Glaze et al, 2015; Veliz-Cuba et al, 2016). Wilson et al (2010) developed a model of an observer that infers the rate of environmental change from observations. To do so, the observer computes a joint posterior probability of the state of the environment, and the time since the last change in the environment. With more measurements, such observers improve their estimates of the change rate, and are therefore better able to predict the environmental state. Inference of the changerate is most important when an observer makes fairly noisy measurements, and cannot determine the current state from a single observation.

We extend previous accumulator models of decision making to the case of multiple, discrete choices with asymmetric, unknown transition rates between them. We assume that the observer is primarily interested in the current state of the environment, often referred to as the *correct choice* in decision-making models (Bogacz et al, 2006). Therefore, we show how an ideal observer can use sensory evidence to infer the rates at which the environment transitions between states, and simultaneously use these inferred rates to discount old evidence and determine the present environmental state.

Related models have been studied before (Wilson et al, 2010; Adams and MacKay, 2007). However, they relied on the assumption that after a change the new state does not depend on the previous state. This excludes the possibility of a finite number of states: For example, with two choices knowledge of the present state determines with complete certainty the state after a change, and the two are thus not independent. In the case of a finite number of choices our algorithm is simpler than previous ones. The observer only needs to compute a joint probability of the environmental state and the number of changepoints.

The storage needed to implement our algorithms grows rapidly with the number of possible environmental states. However, we show that moment closure methods can be used to decrease the needed storage considerably, albeit at the expense of accuracy and the representation of higher order statistics. Nonetheless, when measurement noise is not too large, these approximations can be used to estimate the most likely transition rate, and the current state of the environment. This motivates a physiologically plausible neural implementation for the present computation: We show that a Hebbian learning rule which shapes interactions between multiple neural populations representing the different choices, allows a network to integrate inputs nearly optimally. Our work therefore links statistical principles for optimal inference with stochastic neural rate models that can adapt to the environmental volatility to make near-optimal decisions in a changing environment.

## 2 Optimal evidence accumulation for known transition rates

We start by revisiting the problem of inferring the current state of the environment from a sequence of noisy observations. We assume that the number of states is finite, and the state of the environment changes at times unknown to the observer. We first review the case when the rate of these changes is known to the observer. In later sections we will assume that these rates must also be learned. Following Veliz-Cuba et al (2016), we derived a recursive equation for the likelihoods of the different states, and an approximating stochastic differential equation (SDE). Similar derivations were presented for decisions between two choices by Deneve (2008), and Glaze et al (2015).

An ideal observer decides between *N* choices, based on successive observations at times *t_n_* (*n* = 1, 2,…). We denote each possible choice by *H^i^* (*i* = 1, …, *N*), with *H_n_* being the correct choice at time *t_n_*. The *transition rates ∊^ij^, i* ≠ *j*, correspond to the *known* probabilities that the state changes between two observations: *∊^ij^* = P (*H_n_* = *H^i^*|*H*_*n*−1_ = *H^j^*). The observer makes measurements, *ξ_n_*, at times *t_n_* with known conditional probability densities *f^i^(ξ)* = P (*ξ_n_* = |*H_n_* = *H^i^*). Here, and elsewhere, we assume that the observations are conditionally independent. We also abuse notation slightly by using P(·) to denote a probability, or the value of a probability density function, depending on the argument. We use explicit notation for the probability density function when there is a potential for confusion.

We denote by *ξ_j:n_* the vector of observations (*ξ_j_*,…, *ξ_n_*), and by P_*n*_(·) the conditional probability P(·|*ξ*_1:*n*_). To make a decision, the observer can compute the index that maximizes the *posterior probability*, *î* = argmax_*i*_ P_*n*_(*H_n_* = *H^i^*). Therefore *H*^î^ is the most probable state, given the observations *ξ*_1:*n*_.

A recursive equation for the update of each of the probabilities P_*n*_(*H_n_* = *H^i^*) after the *n*^th^ observation has the form (Veliz-Cuba et al, 2016)

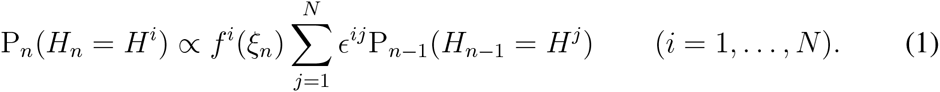

Thus, the transition rates, *∊^jj^*, provide the weights of the previous probabilities in the update equation. Unless transition rates are large or observations very noisy, the probability P_*n*_(*H_n_* = *H^î^*) grows, and can be used to identify the present environmental state. However, with positive transition rates, the posterior probabilities tend to saturate at a value below unity. Strong observational evidence that contradicts an observer’s current belief can cause the observer to change their belief subsequently. Such contradictory evidence typically arrives after a change in the environment.

Taking logarithms, 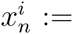 ln *P_n_*(*H_n_ = H^j^*), and assuming that evidence from each observation, as well as the time between observations, Δ*t* := *t_n_ − t_n−1_*, are small, we can approximate the discrete process Eq. (1) with an SDE,

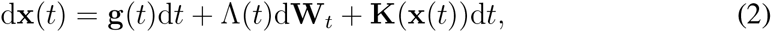

where 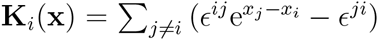, and the drift, 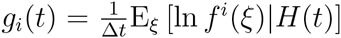, depends on the state of the environment at time *t*, *H(t)* ∈ {*H*^1^,…, *H^N^*}. The drift, *g_i_(t)*, is largest when the environment is in state *H^i^*. The noise covariance, Λ(*t*)Λ(*t*)^*T*^ = Σ(*t*), has entries 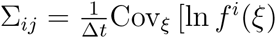, ln *f^j^* (ξ)|*H* (*t*)], while W_*t*_ is a vector of independent Wiener processes.

The nonlinear term, K_*i*_(x), implies that in the absence of noise Eq. (2) has a stable fixed point, and older evidence is discounted. Such continuum models of evidence accumulation are useful because they are amenable to the methods of stochastic analysis (Bogacz et al, 2006). Linearization of the SDE provides insights into the system’s local dynamics (Glaze et al, 2015; Veliz-Cuba et al, 2016), and can be used to implement the inference process in model neural networks (Bogacz et al, 2006; Veliz-Cuba et al, 2016).

We next extend this approach to the case when the observer infers the transition rates, *∊^ij^*, from measurements.

## 3 Environments with symmetric transition rates

We first derive the ideal observer model when the unknown transition rates are symmetric, *∊^ij^* ≡ constant when *j* ≠ *i*, and *∊^ij^* := 1 – (*N* – 1)*∊^ij^*. This simplifies the derivation, since the observer only needs to estimate a single changepoint count. The general, asymmetric case discussed in Section 4 follows the same idea, but the derivation is more involved since the observer must estimate multiple counts.

Our problem differs from previous studies in two key ways (Adams and MacKay, 2007; Wilson et al, 2010): We assume the observer tries to identify the most likely state of the environment at time *t_n_*. To do so the observer computes the joint conditional probability, P_*n*_(*H_n_, a_n_*), of the current state, *H_n_*, and the number of environmental changes since beginning the observation, *a_n_*. Previous studies focused on obtaining the predictive distribution, P_*n*_(*H*_*n*+1_). The two distributions are closely related, as 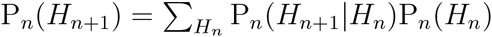.

More importantly, Adams and MacKay (2007); Wilson et al (2010) implicitly assumed that only observations since the last changepoint provide information about the current environmental state. That is, if the time since the last changepoint – the current run-length, *r_n_* – is known to the observer, then all observations before that time can be discarded:

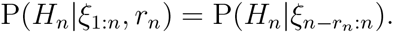

This follows from the assumption that the state after a change is conditionally independent of the state that preceded it. We assume that the number of environmental states is finite. Hence this independence assumption does not hold: Intuitively, if observations prior to a changepoint indicate the true state is *H^j^*, then states *H^i^,i* ≠ *j* are more likely after the changepoint.

Adams and MacKay (2007); Wilson et al (2010) derive a probability update equation for the *run length*, and the number of *changepoints*, and use this equation to obtain the predictive distribution of future observations. We show that it is not necessary to compute run length probabilities when the number of environmental states is finite. Instead we derive a recursive equation for the joint probability of the current state, *H_n_*, and number of changepoints, *a_n_*. As a result, the total number of possible states grows as *N · n* (linearly in *n*) where *N* is the fixed number of environmental states *H^i^* rather than *n*^2^ (quadratically in *n*) as in Wilson et al (2010).

### 3.1 Symmetric 2-state process

We first derive a recursive equation for the probability of two alternatives, *H_n_* ∈ {*H*^±^}, in a changing environment, where the change process is memoryless, and the change rate, 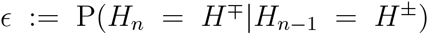, is symmetric and initially unknown to the observer (See Fig. 1A). The most probable choice given the observations up to a time, *t_n_*, can be obtained from the log of the posterior odds ratio 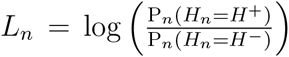. The sign of *L_n_* indicates which option is more likely, and its magnitude indicates the strength of this evidence (Bogacz et al, 2006; Gold and Shadlen, 2007). Old evidence should be discounted according to the inferred environmental volatility. Since this is unknown, an ideal observer computes the probability distribution of the change rate, *∊* (See Fig. 1C), along with the probability of environmental states.

**Figure 1:**
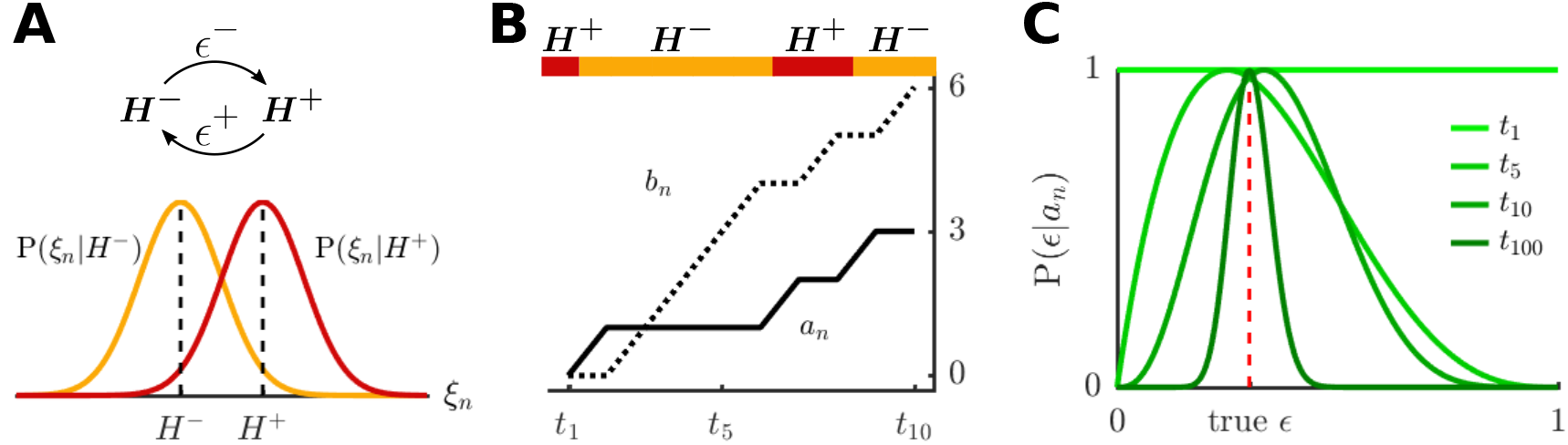
Online inference of the change rate in a dynamic environment. (A) The environment alternates between states *H*^+^ and *H*^−^ with transition probabilities *∊*+, *∊*^−^. The state of the environment determines *f*^±^(*ξ*) = P(*ξ*|*H*^±^), which we represent as Gaussian densities. (B) A sample path of the environment (color bar) together with the first ten values of the actual changepoint count, *a_n_*, and non-changepoint count, *b_n_*. (C) Evolution of the conditional probabilities, P(*∊*|*a_n_*), corresponding to the changepoint count from panel B, until *t_n_* = *t*_100_. The dashed red line indicates the value of *∊* in the simulation. The densities are scaled so that each equals 1 at the mode.

Let *a_n_* be the number of changepoints, and *b_n_* = *n* – 1 – *a_n_* the count of non-changepoints between times t_1_ and *t_n_* (*n* = 1,2,…) (See Fig. 1B). The process {*a_n_*}_*n*≥1_ is a pure birth process with birth rate *∊*. The observer assumes no changes prior to the start of observation, P(*a*_1_ = 0) = 1, and must make at least two observations, *ξ*_1_ and *ξ*_2_, to detect a change.

To develop an iterative equation for the joint conditional probability density, P_*n*_(*H_n_, a_n_*), given the n observations *ξ*_1:*n*_, we begin by marginalizing over these quantities at the time of the previous observation, *t*_*n*−1_, for *n* > 1 (See Appendix 7.1 for details):

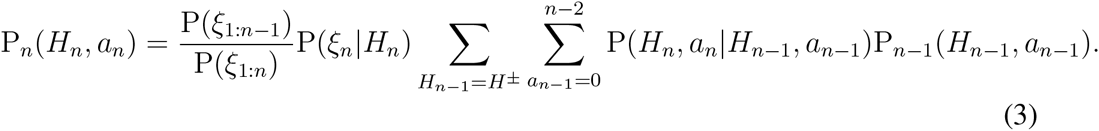

With two choices we have the following relationships for all n > 1:

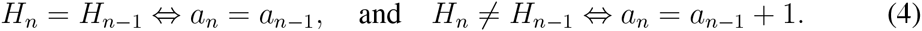

The term P(*H_n_, a_n_*|*H_n−1_, a_n−1_*) in Eq. (3) is therefore nonzero only if either, *H_n−1_* = *H_n_*, and *a_n−1_ = a_n_*, or *H_n−1_ = H_n_* and *a_n−1_ = a_n_* – 1: If the system is in the joint state (*H_n−1_, a_n−1_*) at *t_n−1_*, then at *t_n_* it can either (a) transition to (*H_n_ = H_n−1_, a_n_ = a_n−1_* + 1) or (b) remain at (*H_n_ = H_n−1_, a_n_ = a_n−1_*). This observation is central to the message-passing algorithm described in (Adams and MacKay, 2007; Wilson et al, 2010), with probability mass flowing from lower to higher values of a according to a pure birth process (See Fig. 2A). We can thus simplify Eq. (3), leaving only two terms in the double sum. Writing P_*n*_ (*H^±^, a*) for P_*n*_ (*H_n_ = H^±^, a_n_ = a*), and similarly for any conditional probabilities, we have for *n* > 1:

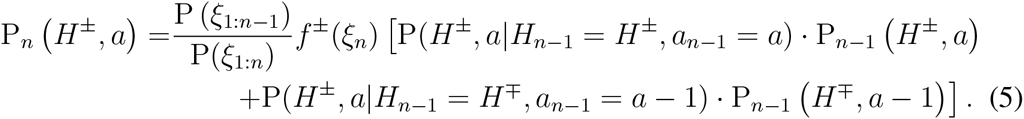

**Figure 2:**
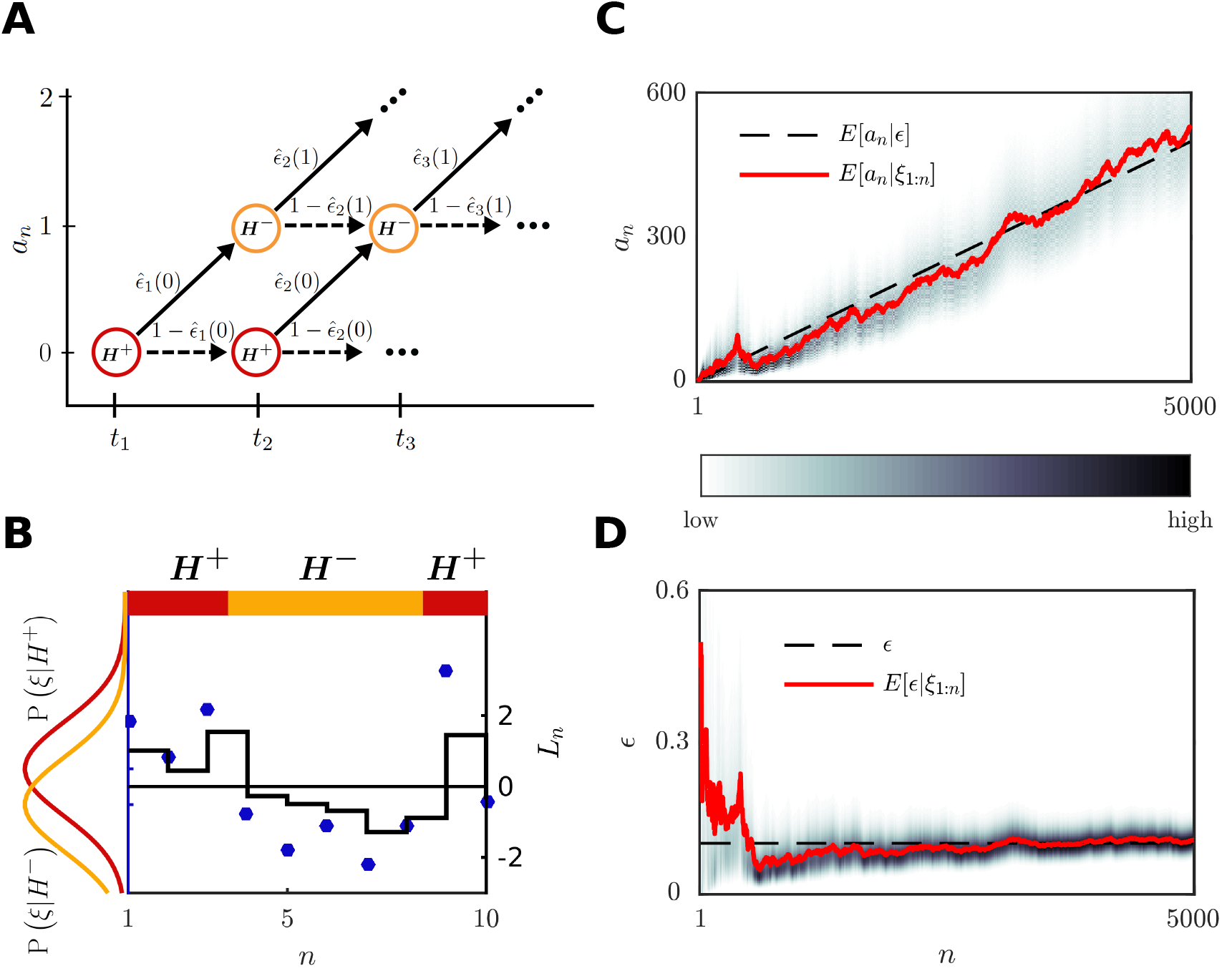
Inference of the state, *H_i_*, and change rate, *∊*. (A) The joint posterior probability, P_*n*_(*H^±^, a*), is propagated along a directed graph according to Eq. (17). Only paths corresponding to the initial condition (*H*_1_, *a*_1_) = (*H*^+^, 0) are shown. (B) A sample sequence of environmental states (color bar, top) together with the first ten observations *ξ*_1_,… *ξ*_10_ (blue dots), for *∊*^+^ = *∊*^−^ = 0.1. Superimposed in black (right y-axis) is the log posterior odds ratio *L_n_* as a function of time. (C) Evolution of the posterior over *a_n_* (gray scale). The posterior mean (red) converges to the expected number of changepoints *∊*(*n* – 1) (dashed line). (D) Evolution of the posterior over the change rate *∊* (gray scale). The posterior mean (red) converges to the true value (dashed line) and the variance diminishes with the number of observations.

We must also specify *initial conditions* at time *t*_1_, and *boundary values* when *a* ∈ {0, *n* – 1} for these equations. At *t*_1_ we have P(*a*_1_ = 0) = 1. Therefore,

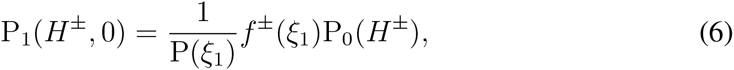

and P_1_(*H^±^, a*) = 0 for *a* = 0. Here P_0_(*H^±^*) is the prior over the two choices. The probability P(*ξ*_1_) is unknown to the observer. However, similar to the ratio 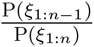 in Eq. (5), P(*ξ*_1_) acts as a normalization constant and does not appear in the posterior oddsratio, *R_n_* (See Eq. (18) below). Finally, at all future times *n* > 1, we have separate equations at the boundaries,

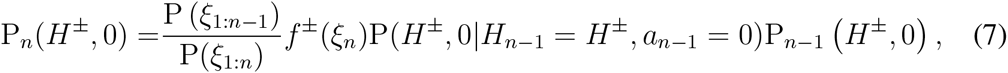

and,

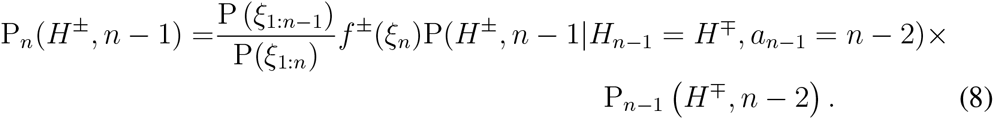

We next compute P(*H_n_, a_n_|H_n−1_, a_n−1_*) in Eq. (3), with *n* > 1, by marginalizing over all possible transition rates ∊ ∈ [0, 1]:

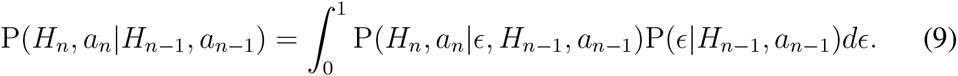

Note that P(*∊|H_n−1_, a_n−1_*) = P(*∊*|*a_n−1_*), so we need the distribution of *∊*, given *a_n−1_* changepoints, for all *n* > 1. We assume that prior to any changepoint observations — that is at time *t*_1_ — the rates follow a Beta distribution with hyperparameters *a*_0_, *b*_0_ > 0 (See also Sections 3.1 and 3.2 in Wilson et al (2010)),

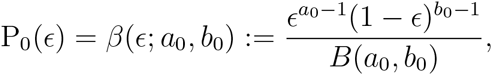

where *β* denotes the probability density of the associated Beta distribution, and 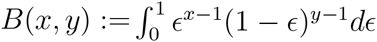 is the beta function. For any *n* > 1, the random variable *a_n_*|*∊* follows a Binomial distribution with parameters (*n* – 1, *∊*), for which the Beta distribution is a conjugate prior. The posterior over the change rate when the changepoint count is known at time *n* > 1 is therefore:

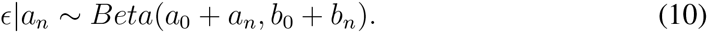

For simplicity, we assume that prior to any observations, the probability over the transition rates is uniform, P_0_(*∊*) = 1 for all *∊* ∈ [0,1], and therefore *a*_0_ = *b*_0_ = 1.

We now return to Eq. (9) and use the definition of the transition rate, *∊*, (See Fig. 1) to find:

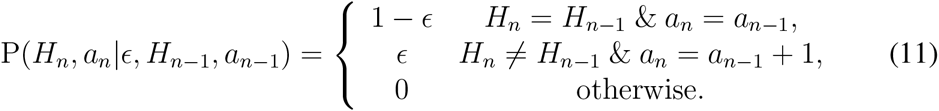

Eq. (9) can therefore be rewritten using two integrals, depending on the values of (*H_n_, a_n_*) and (*H_n−1_, a_n−1_*),

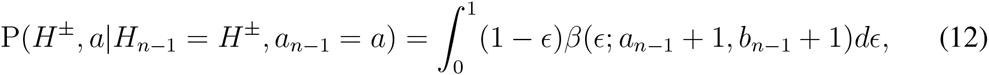

and similarly for 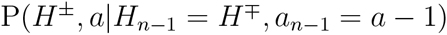.

The mean of the Beta distribution, for *n* > 1, can be expressed in terms of its two parameters:

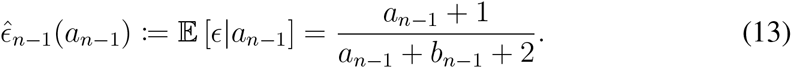

We denote this expected value by 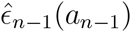 as it represents a point estimate of the change rate *∊* at time *t_n−1_* when the changepoint count is *a_n−1_, n* > 1. Since *a_n−1_* + *b_n−1_ = n* – 2, we have:

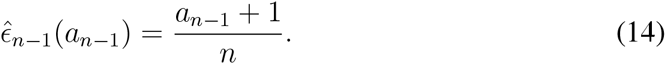

The expected transition rate, 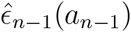, is thus determined by the ratio between the present changepoint count and the number of timesteps, *n*. Leaving *a*_0_ and *b*_0_ as parameters in the prior gives 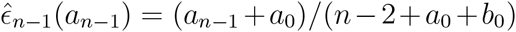. Using the definition in Eq. (14), it follows from Eq. (12) that:

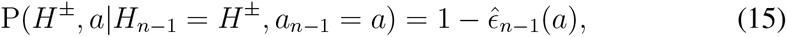

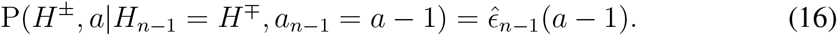

Eqs. (15) and (16), which are illustrated in Fig. 2A, can in turn be substituted into Eq. (5) to yield, for all *n* > 1:

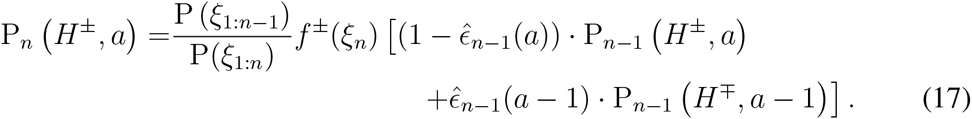

The initial conditions and boundary equations for this recursive probability update have already been described in Eqs. (6–8). Eq. (17) is the equivalent of Eq. (3) in Adams and MacKay (2007), and Eq. (3.7) in Wilson et al (2010). However, here the observer does not need to estimate the length of the interval since the last changepoint. We demonstrate the inference process defined by Eq. (17) in Fig. 2.

The observer can compute the posterior odds ratio by marginalizing over the change-point count:

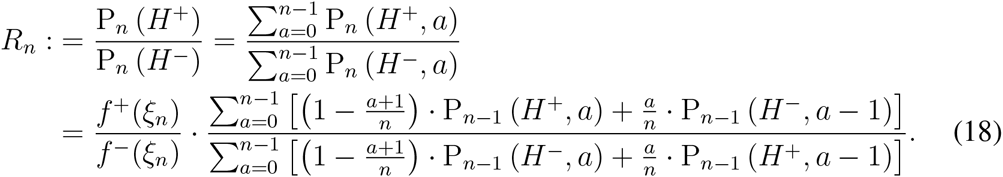

Here log(*R_n_*) = *L_n_* > 0 implies that *H_n_* = *H^+^* is more likely than *H_n_* = *H^−^* (See Fig. 2B). Note that P(*ξ*_1:*n*−1_)|P(*ξ*_1:*n*_) and 1/P(*ξ*_1_) need not be known to the observer to obtain the most likely choice.

A posterior distribution of the transition rate e can also be derived from Eq. (17) by marginalizing over (*H_n_, a_n_*),

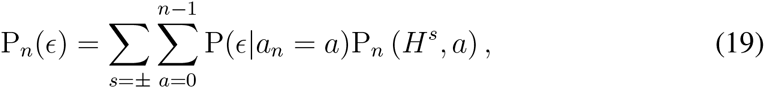

where P(*∊|a_n_*) is given by the Beta distribution prior Eq. (10). The expected rate is therefore:

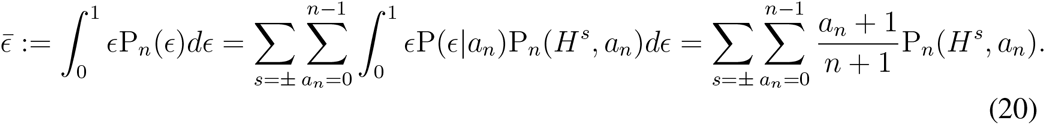

Explicit knowledge of the transition rate, *∊*, is not used in the inference process described by Eq. (17). However, computing it allows us to evaluate how the observer’s estimate converges to the true transition rate (See Fig. 2D). We will also relate this estimate to the coupling strength between neural populations in the model described in Section 6.

We conjecture that when measurements are noisy, the variance of the distribution P_*n*_(*∊*) does not converge to a point mass at the true rate, *∊*, in the limit of infinitely many observations, *n* → ∞, *i.e*. the estimate of *∊* is not consistent. As we have shown, to infer the rate we need to infer the parameter of a Bernoulli variable. It is easy to show that the posterior over this parameter converges to a point mass at the actual rate value if the probability of misclassifying the state is known to the observer (Djuric and Huang, 2000). However, when the misclassification probability is not known, the variance of the posterior remains positive even in the limit of infinitely many observations. In our case, when measurements are noisy, the observer does not know the exact number of change points at finite time. Hence, the observer does not know exactly how to weight previous observations to make an inference about the current state. As a result, the probability of misclassifying the current state may not be known. We conjecture that this implies that even in the limit *n* → ∞ the posterior over *∊* has positive variance (See Fig. 2D).

In Fig. 3 we compare the performance of this algorithm in three cases: when the observer knows the true rate (point mass prior over the true rate *∊*); when the observer assumes a wrong rate (point mass prior over an erroneous *∊*); and when the observer learns the rate from measurements (flat prior over *∊*). We define performance as the probability of a correct decision.

**Figure 3:**
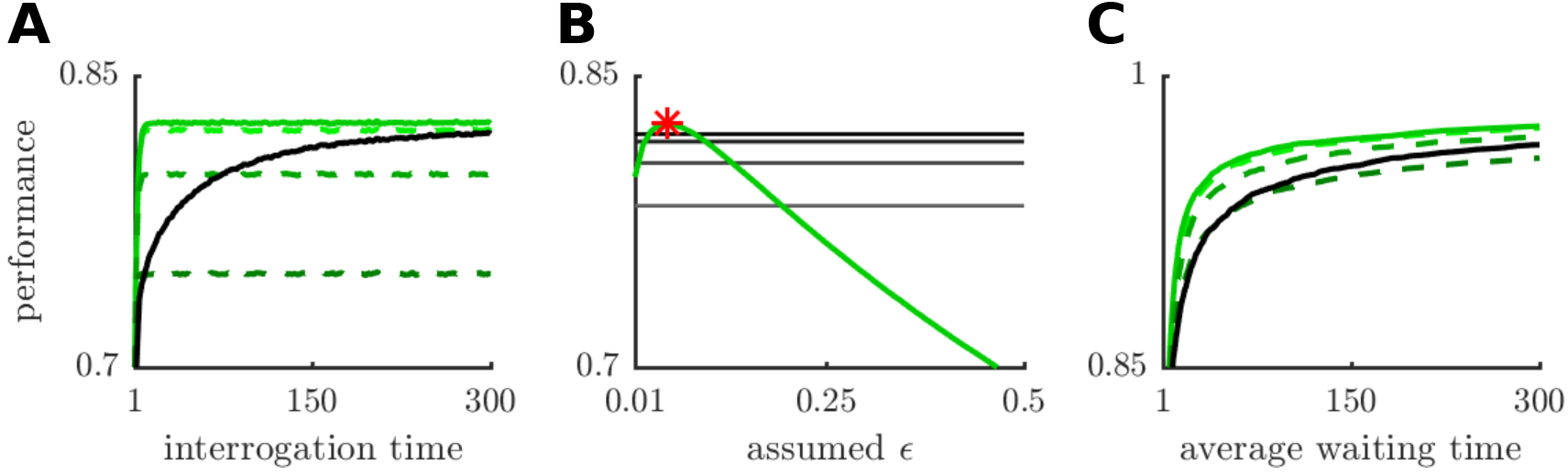
The performance of the inference algorithm. (A) Performance under the interrogation paradigm measured as the percentage of correct responses at the interrogation time. Here and in the next panel *∊* = 0.05, and SNR= 1. The black curve represents the performance of an ideal observer who infers the change rate from measurements. The green curves, represent the performance of observers that assume a fixed change rate (0.3, 0.15, 0.05, 0.03 from darker to lighter, see Eq. (1)). The solid green line corresponds to the true rate, dashed lines to erroneous rates. (B) The green curve represents the performance at interrogation time *t*_300_ of an observer that assumes a fixed change rate. The red star marks the maximum of this curve. The horizontal black curves represent the performance at times *t*_40_, *t*_100_, *t*_200_, *t*_300_ (from bottom to top) of the observer that learns the change rate. (C) The accuracy as a function of the average threshold hitting time in the free response protocol. Here *∊* = 0.1, and SNR=0.75. See Appendix 7.2 for details on numerical simulations. See also Fig. 3 in Veliz-Cuba et al (2016).

Under the interrogation protocol, the observer infers the state of the environment at a fixed time. As expected, performance increases with interrogation time, and is highest if the observer uses the true rate (See Fig. 3A, also Eq. (1) above). Performance plateaus quickly when the observer assumes a fixed rate, and more slowly if the rate is learned. The performance of observers that learn the rate slowly increases toward that of observers who know the true rate. In panel B, we present the performance of the unknown-rate algorithm at 4 different times (*t*_40_, *t*_100_, *t*_200_, *t*_300_) and compare it to the asymptotic values with different assumed rates (green curves).

Note, an observer that assumes an incorrect change rate can still perform near optimally (e.g., curve for 0.03 in Fig. 3A), especially when the signal-to-noise ratio (SNR) is quite high. The SNR is the difference in means of the likelihoods divided by their common standard deviation. Change rate inference is more effective at lower SNR values, in which case multiple observations are needed for an accurate estimate of the present state. However, at very low SNR values the observer will not be able to substantially reduce uncertainty about the change rate, resulting in high uncertainty about the state.

In the free response protocol, the observer makes a decision when the log odds ratio reaches a predefined threshold. In Fig. 3C, we present simulation results for this protocol in a format similar to Fig. 3A, with empirical performance as a function of average hitting time. Each performance level corresponds to unique log odds threshold. Similar to the interrogation protocol (Fig. 3A), performance of the free response protocol saturates much more quickly for an observer that fixes their change rate estimate than on that infers this rate over time.

### 3.2 Symmetric multistate process

We next consider evidence accumulation in an environment with an arbitrary number of states, {*H*^1^,*H*^2^,…,*H^N^*}, with symmetric transition probabilities, *∊^jj^* constant, whenever *i* ≠ *j*. We define *∊* := (*N* – 1)*∊^ij^* for any *i* ≠ *j*, so that the probability of remaining in the same state becomes *∊^ii^* = 1 – *∊*, for all *i* = 1,…, *N*. The symmetry in transition rates means that an observer still only needs to track the total number of changepoints, *a_n_*, as in Section 3.1.

Eqs. (3–4) remain valid with N possible choices, {*H*^1^,…, *H^N^*}. When *n* > 1, the double sum in Eq. (3) simplifies to:

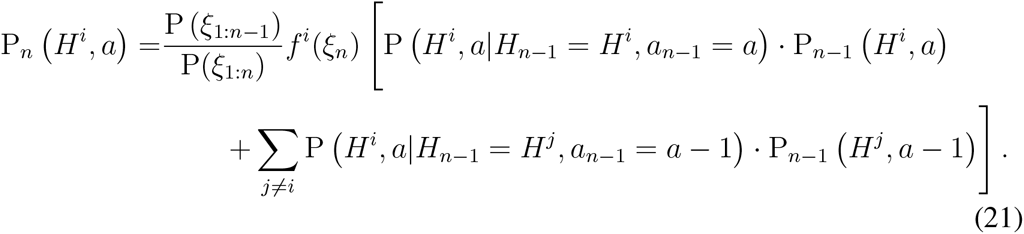

As in Section 3.1, we have P_1_(*H^i^*, 0) = *f^i^*(*ξ*_1_)P_0_(*H^i^*)|P(*ξ*_1_) and P_1_(*H^i^,a*_1_) = 0 for *a*_1_ ≠ 0, where P_0_(*H^i^*) describes the observer’s belief prior to any observations. At all future times, *n* > 1, we have at the boundaries for all *i* = 1,…, *N*:

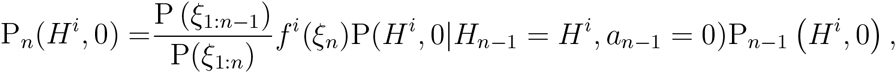

and,

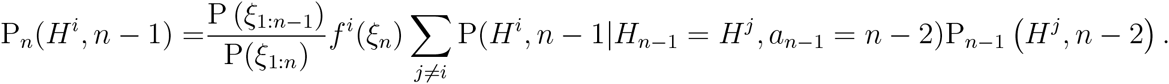

Eq. (9) remains unchanged and we still have P(*∊*|*H_n−1_, a_n−1_*) = P(*∊|a_n−1_*). Furthermore, assuming a Beta prior on the change rate, Eq. (10) remains valid, and Eq. (11) is replaced by:

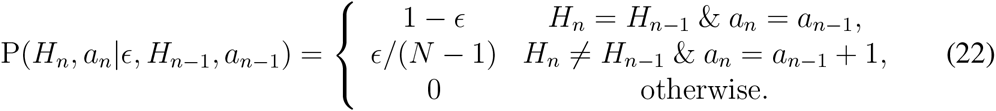

The integral from Eq. (9) gives, once again, the mean of the Beta distribution, 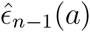, defined in Eqs. (13–14). As in Section 3.1, 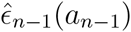 is a point estimate of the change rate *∊* at time *t_n−1_* when the changepoint count is *a_n−1_*. We have,

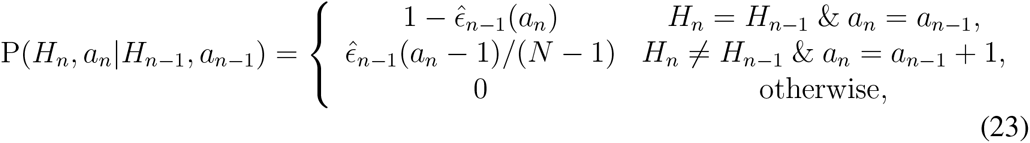

and the main probability update equation is now:

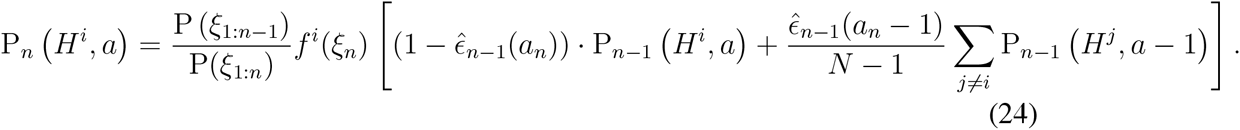

The observer can infer the most likely state of the environments, by computing the index that maximizes the *posterior probability*, marginalizing over all changepoint counts,

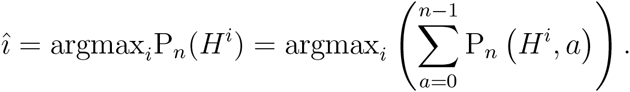

The observer can also compute the posterior probability P_*n*_(*∊*) of the transition rate, *∊*, by marginalizing over all states *H_n_* and changepoint counts *a_n_*, as in Eq. (19). Furthermore, a point estimate of ∊ is given by the mean of the posterior after marginalizing, as in Eq. (20).

## 4 Environments with asymmetric transition rates

In this section, we depart from the framework of Adams and MacKay (2007), and Wilson et al (2010), and consider unequal transition rates between states. This includes the possibility that some transitions are not allowed. We consider an arbitrary number, *N*, of states with unknown transition rates, *∊^ij^*, between them. The switching process between the states is again memoryless, so that *H_n_* is a stationary, discrete-time Markov chain with finite state space, Ω := {*H*^1^,…, *H^N^*}. We write the (unknown) transition matrix for this chain as a left stochastic matrix,

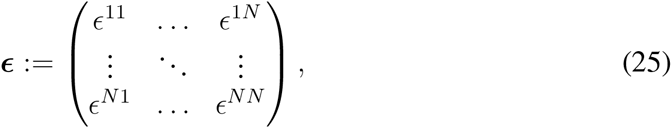

where *∊^ij^* = P(*H_n_ = H^i^|H_n−1_ = H^j^*), with *i, j* ∈ {1,…, *N*}. We denote by *∊^.i^* the *i*-th column of the matrix *∊*, and similarly for other matrices. Each such column sums to 1. We define the changepoint counts matrix at time *t_n_* as,

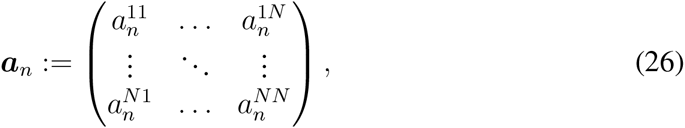

where 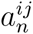 is the number of transitions from state *j* to state *i* up to time *t_n_*. There can be a maximum of *n* – 1 transitions at time *t_n_*. For a fixed *n* ≥ 1, all entries in *a_n_* are nonnegative and sum to *n* – 1, *i.e*. 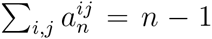. As in the symmetric case, the changepoint matrix at time *t*_1_ must be the zero matrix, *a*_1_ = 0.

We will show that our inference algorithm assigns positive probability only to changepoint matrices that correspond to possible transition paths between the states {*H*^1^,…, *H^N^*}. Many nonnegative integer matrices with entries that sum to *n* – 1 are not possible changepoint matrices *a_n_*. A combinatorial argument shows that when *N* = 2 state case scales as *n*^2^, the number of possible pairs, (*H_n_, a_n_*), grows quadratically with the number of steps, *n*, to leading order. It can also be shown that the growth is polynomial for *N* > 2, although we do not know the growth rate in general (See Fig. 4B). An ideal observer has to assign a probability of each of these states which is much more demanding than in the symmetric rate case where the number of possible states grows linearly.

**Figure 4:**
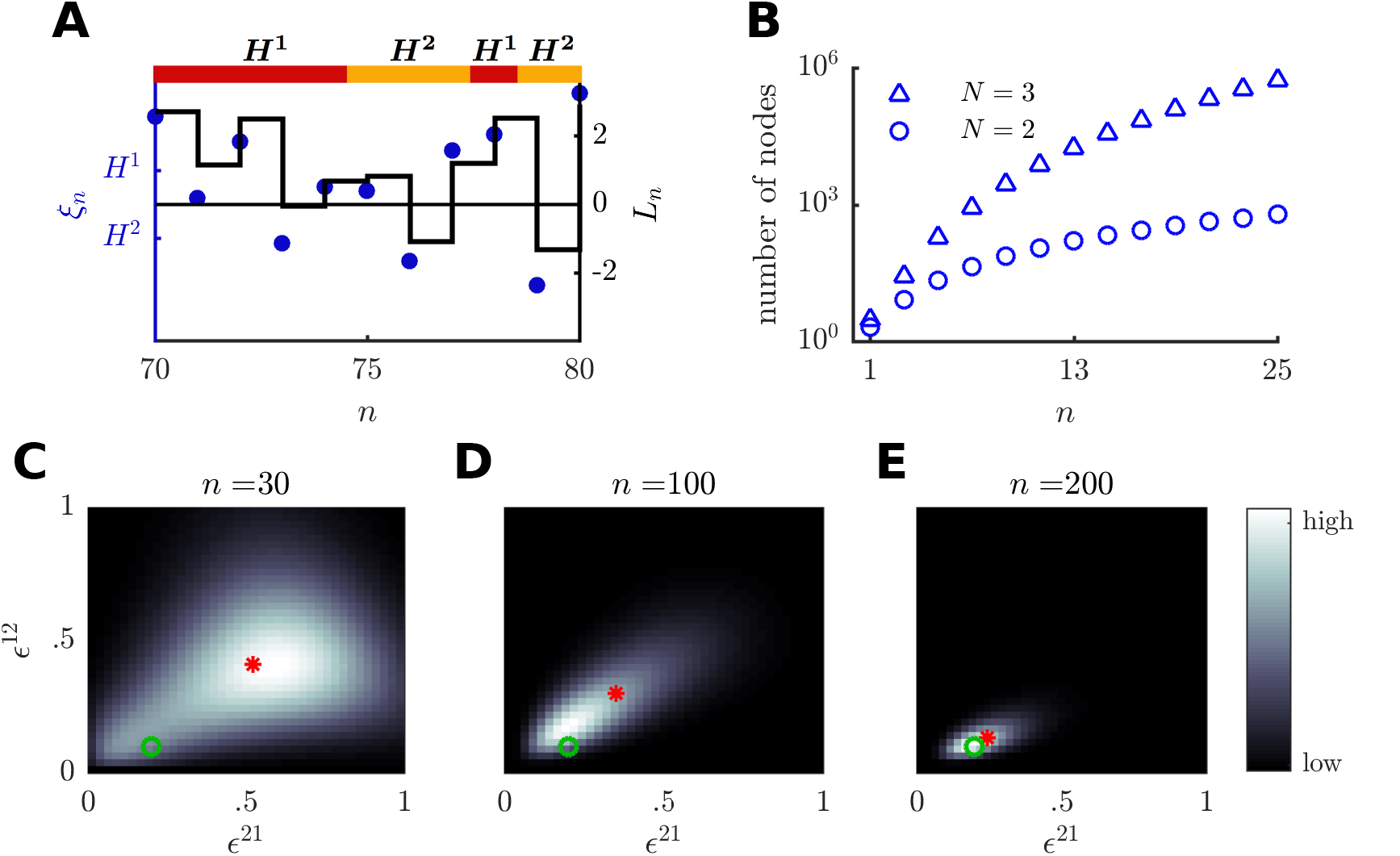
Evidence accumulation and change rates inference in a 2-state asymmetric system. (A) Sample path (color bar, top) of the environment between times *t*_70_ and *t*_80_ (same simulation as in panels C-E) with corresponding observations (blue dots), and log posterior odds ratio (black step function). Here and in panels C-E, (*∊*^+^, *∊*^−^) = (0.2,0.1), SNR= 1.4. (B) The number of allowable changepoint matrices as a function of observation number, *n*, for *N* = 2 (blue circles), and *N* = 3 (blue triangles). (C)-(E) Color plots (gray scale) of the joint density, P_*n*_ (*∊*^21^, *∊*^12^), with mean value (red star) approaching the true transition rates (green circle).

We next derive an iterative equation for P_*n*_(*H_n_, a_n_*), the joint probability of the state *H_n_*, and an allowable combination of the *N*(*N* – 1) changepoint counts (off-diagonal terms of *a_n_*), and *N* non-changepoint counts (diagonal terms of *a_n_*). The derivation is similar to the symmetric case: For *n* > 1, we first marginalize over *H_n−1_* and *a_n−1_*,

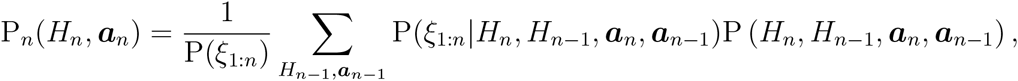

where the sum is over all *H_n−1_* ∈ {*H*^1^,…, *H^N^*} and possible values of the changepoint matrix, *a_n−1_*.

Using P(*H_n_*, *H_n−1_, a_n_, a_n−1_*) = P(*H_n_, a_n_|H_n−1_, a_n−1_*)P(*H_n−1_, a_n−1_*), and applying Bayes’ rule to write

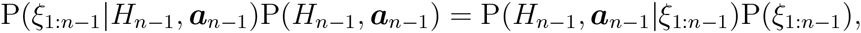

gives

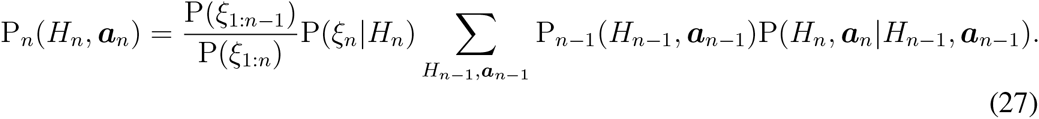

We compute the conditional probability P(*H_n_, a_n_|H_n−1_, a_n−1_*) by marginalizing over all possible transition matrices, *∊*. To do so, we relate the probabilities of *∊* and *a*. Note that if the observer assumes the columns *∊^.j^* are independent prior to any observations, then the exit rates conditioned on the changepoint counts, 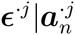, are independent for all states, *j* = 1,…, *N*.

To motivate the derivation we first consider a single state, *j* = 1, and assume that the environmental state has been observed perfectly over *T* > 1 timesteps, but the transition rates are unknown. Therefore, all 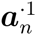 are known to the observer (1 ≤ *n* ≤ *T*), but the *∊*^·1^ are not. The state of the system at time *n* + 1, given that it was in state *H*^1^ at time *n*, is a categorical random variable, and P(*H_n+1_* = *H^i^|H_n_ = H*^1^) = *∊*^*i*1^, for 1 ≤ *n* ≤ *T* – 1. The observed transitions *H*^1^ ↦ *H^i^* are independent samples from a categorical distribution with unknown parameters *∊*^·1^.

The conjugate prior to the categorical distribution is the Dirichlet distribution, and we therefore use it as a prior on the changepoint probabilities. For simplicity we again assume a flat prior over *∊*^·1^, that is P(*∊*^·1^) = χ_*S*_(*∊*^·1^), where χ_*S*_ is the indicator function on the standard (*N* – 1)-simplex, *S*.

Denote by *D* the sequence of states that the environment transitioned to at time *n* + 1 whenever it was in state *H*^1^ at time *n*, for all 1 ≤ *n* ≤ *T* – 1. Therefore *D* is a sequence of states from the set {*H*^1^,…, *H^N^*}. By definition, 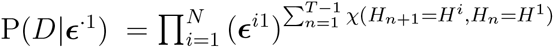, where χ(*H_n+1_* = *H^i^, H_n_* = *H*^1^) is the indicator function, which is unity only when *H_n+1_* = *H^i^* and *H_n_* = *H*^1^ and zero otherwise. Equivalently, we can write 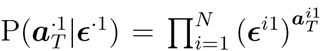, since 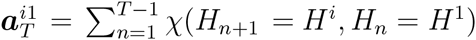. For general *n* > 1, the posterior distribution for the transition probabilities *∊*^·1^ given the changepoint vector 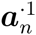 is then

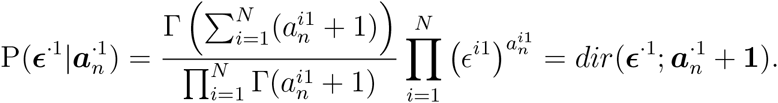

Here 1 = (1,…, 1)^*T*^, so 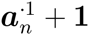 should be interpreted as the vector with entries 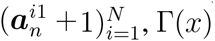 is the gamma function, and 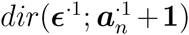 the probability density function of the *N*-dimensional Dirichlet distribution, 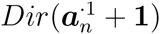.

The same argument applies to all initial states, *H^j^, j* ∈ {1,…, *N*}. We assumed that the transition rates are conditionally independent, so that

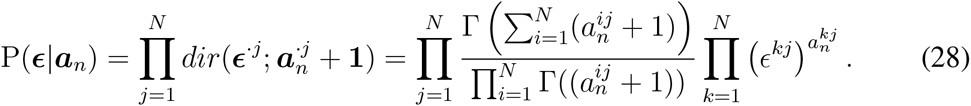

Using this observation, the transition probability between two states can be computed by marginalizing over all possible transition matrices, *∊*, conditioned on *a_n–1_*,

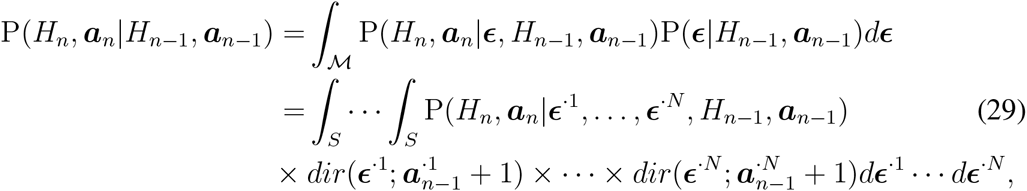

where 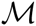 represents the space of all *N × N* left stochastic matrices and *S* is the *N* – 1 dimensional simplex of *∊*^·*j*^ ∈ [0,1]^*N*^ such that 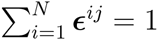.

Let *δ^ij^* be the *N x N* matrix containing a 1 as its *ij*-th entry, and 0 everywhere else. For all *i, j* ∈ {1,…, *N*} we have

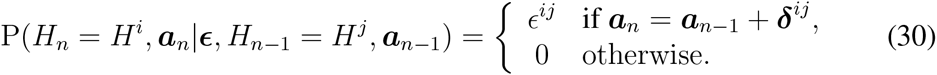

Implicit in Eq. (30) is the requirement that the environment must have been in the state *H_n−1_ = H^j^* in order for the transition *H^j^* ↦ *H^i^* to have occurred between *t_n−1_* and *t_n_*. This will ensure that the changepoint matrices *a_n_* that are assigned nonzero probability correspond to admissible paths through the states {*H*^1^,…, *H^N^*}. Applying Eq. (30), we can compute the integrals in Eq. (29) for all pairs (*i, j*). We let 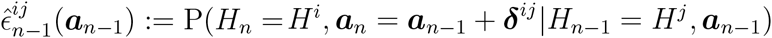 to simplify notation, and find

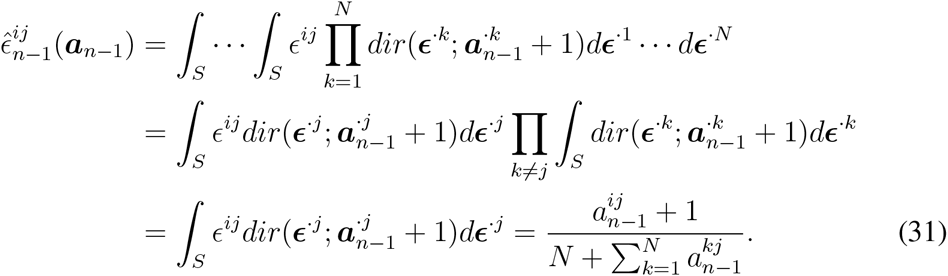

As in the point estimate of the rate 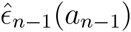 in Eq. (14), each 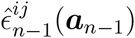 is a ratio containing the number of *H^j^* ↦ *H^i^* transitions in the numerator, and the total number of transitions out of the *j*th state in the denominator. Thus, the estimated transition rate 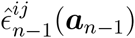 increases with the number of transitions *H^j^* ↦ *H^i^* in a given interval {1,…, *n*}. Furthermore, each column sums to unity:

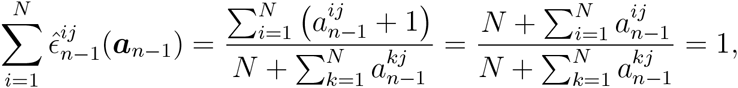

so the point estimates 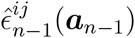 for the transition rates out of each state *j* do provide an empirical probability mass function along each column. However, as in the symmetric case, these estimates are biased toward the interior of the domain. This is a consequence of the hyperparameters we have chosen for our prior density, dir(*∊; a*_0_ + 1).

Therefore, for *n* > 1, the probability update equation in the case of asymmetric transition rates (Eq. (27)) is given by,

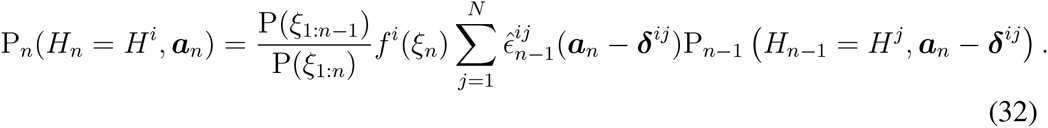

The point estimates of the transition rates, 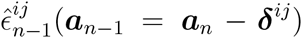, are defined in Eq. (31). As before, 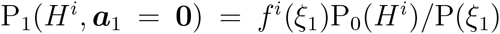 and P_1_(*H^i^*|*a*_1_) = 0 for any *a*_1_ ≠ 0. At future times, it is only possible to obtain changepoint matrices *a_n_* whose entries sum to 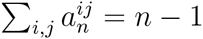, the changepoint matrices *a_n_* and *a_n−1_* must be related as *a_n_* = *a_n−1_* + *δ^ij^*, as noted in Eq. (30). This considerably reduces the number of terms in the sum in Eq. (32).

The observer can find the most likely state of the environment by maximizing the posterior probability after marginalizing over the changepoint counts *a_n_*,

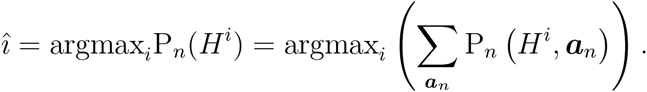

The transition rate matrix can also be computed by marginalizing across all possible states, *H_n_*, and changepoint count matrices, *a_n_*,

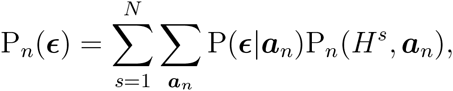

where P(*∊*|*a_n_*) is the product of probability density functions, 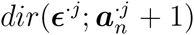, given in Eq. (28). The mean of this distribution is given by

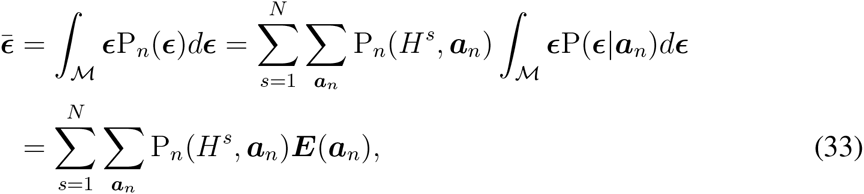

where 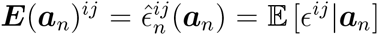 defined in Eq. (31), is a conditional expectation over each possible changepoint matrix *a_n_*.

Eq. (32) is easier to interpret when *N* = 2. Using Eq. (31), we find

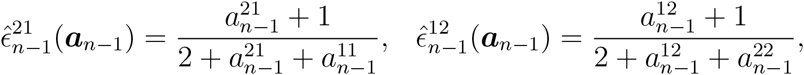

and we can express 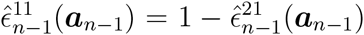 and 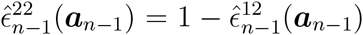. Expanding the sum in Eq. (32), we have

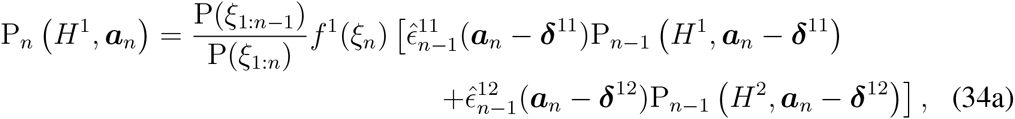

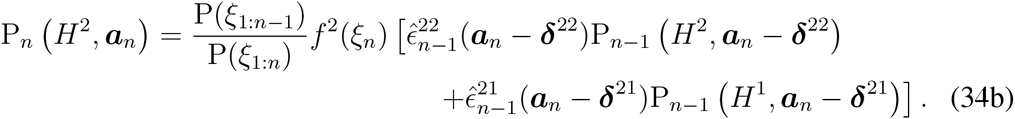

The boundary and initial conditions will be given as above, and the mean inferred transition matrix is given by Eq. (33). Importantly, the inference process described by Eqs. (34) allows for both asymmetric changepoint matrices, *a_n_*, and inferred transition rate matrices ***E**(a_n_)*, unlike the process in Eq. (17). However, the variance of the posteriors over the rates will decrease more slowly, as fewer transitions out of each particular state will be observed.

This algorithm can be used to infer unequal transition rates as shown in Fig. 4: Panels C through E show that the mode of the joint posterior distribution, P_*n*_(*∊*^21^, *∊*^12^), approaches the correct rates, while its variance decreases. As in Section 3.1 we conjecture that this joint density does not converge to a point mass at the true rate values unless the SNR is infinite.

## 5 Continuum limits and stochastic differential equation models

We next derive continuum limits of the discrete probability update equations for the symmetric case discussed in Section 3. We assume that observers make measurements rapidly, so we can derive a stochastic differential equation (SDE) that models the update of an ideal observer’s belief (Gold and Shadlen, 2007). SDEs are generally easier to analyze than their discrete counterparts (Gardiner, 2004). For example, response times can be studied by examining mean first passage times of log-likelihood ratios (Bogacz et al, 2006), or log-likelihoods (McMillen and Holmes, 2006), which is much easier done in the continuum limit (Redner, 2001). For simplicity, we begin with an analysis of the two state process, and then extend our results to the multistate case.

### 5.1 Derivation of the continuum limit

Two-state symmetric process. We first assume that the state of the environment, {*H_t_*}, is a homogeneous *continuous-time* Markov chain with state space {*H*^+^,*H*^−^}. The probability of transitions between the two states is symmetric, and given by 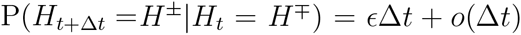, where 0 ≤ *∊* < ∈. The number of changepoints, *a_t_*, up to time *t* is a Poisson process with rate *∊*. An observer infers the present state from a sequence of observations, *ξ*_1:*n*_, made at equally spaced times, *t*_1:*n*_, with Δ*t* = *t_j_* — *t_j−1_*.^1^ Each observation, *ξ_n_*, has probability 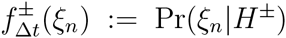 (See Veliz-Cuba et al (2016) for more details). We again use the notation P_*n*_(*H*^±^, *a*) = P(*H_t_n__* = *H*^±^, *a_t_n__* = *a*|*ξ*_1:*n*_) where *t_n_* is the time of the *n*^th^ observation.

As in the previous sections, an estimate of the rate parameter, *∊*, is obtained from the posterior distribution over the changepoint count, *a_t_n__*, at the time of the *n*^th^ observation, *t_n_*. For simplicity, we assume a Gamma prior with parameters *α* and *β* over *∊*, so that *∊* ~ *Gamma*(*α,β*). By assumption the changepoint count follows a Poisson distribution with parameter *∊t_n_*, so that 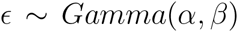. Therefore, once *a_n_* changepoints have been observed, we have the posterior distribution *∊*|*a_n_* ~ Gamma(*a_n_ + a, t_n_ + β*), that is,

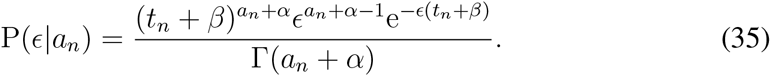

We can substitute Eq. (35) into Eq. (9) describing the probability of transitions between time *t_n−1_* and *t_n_* to find

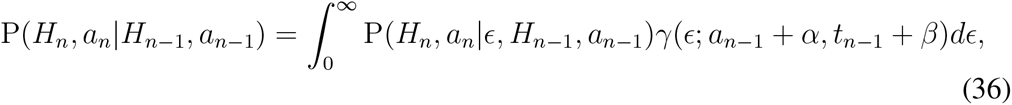

where 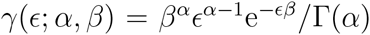 is the density of the Gamma distribution. Using the definition of the transition rate *∊*, we can relate it to the first conditional probability in the integral of Eq. (36) via

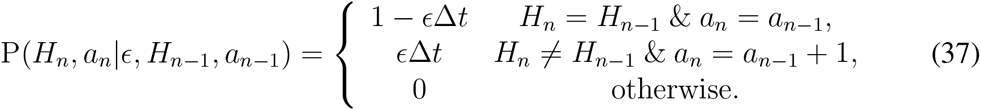

We have dropped the *o*(Δ*t*) terms as we are interested in the limit Δ*t* → 0.

Using Eq. (37) and properties of the Gamma distribution we can evaluate the integral in Eq. (36) to obtain

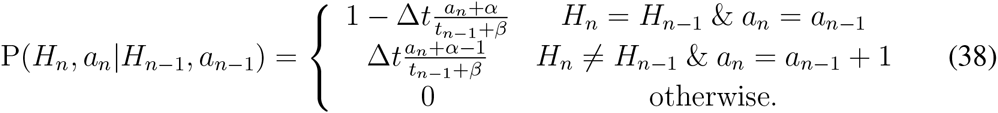

We can use Eq. (38) in the update equation, Eq. (3), to obtain the probabilities of (*H_n_, a_n_*) given observations *ξ*_1:*n*_. As before, only terms involving *a_n_* – 1 and *a_n_* remain in the sum for *n* ≥ 1. Using the same notational convention as in previous sections, we obtain,

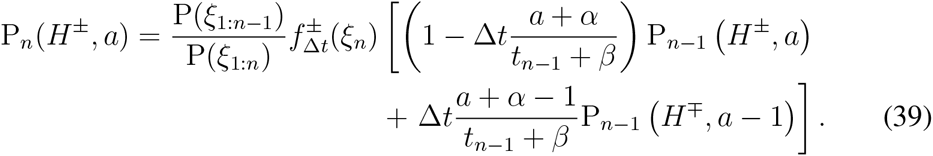

Note, Eq. (39) is similar to the update Eq. (17) we derived in Section 3, with the time index replaced by *t*_*n*−1_/Δ*t* up to the *β* term. Also, since we have used a Gamma instead of a Beta distribution as a prior, the point estimate of the transition rate is slightly different (See Eq. (14)). As in the discrete time case, a point estimate of the transition rate is required even before the first changepoint can be observed. We therefore cannot use an improper prior, as the rate point estimate would be undefined.

To take the limit of Eq. (39) as Δ*t* → 0 we proceed as in Bogacz et al (2006) and Veliz-Cuba et al (2016), working with logarithms of the probabilities. Dividing Eq. (39) by P_*n*−1_ (*H*^±^,*a*), taking logarithms of both sides, and using the notation 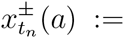 ln P_*n*_(*H*^±^, *a*), we obtain, ^2^

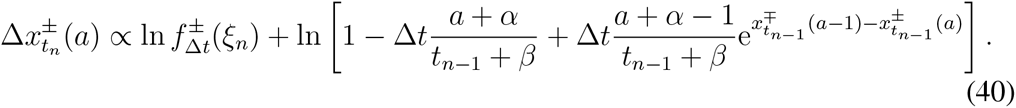

Using the approximation ln(1 + *z*) ≈ *z* for small *z* yields

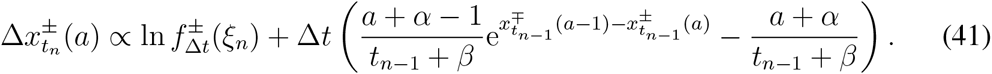

Since the proportionality constant is equal for all *a*, we can use the SDE for the log likelihood *x_t_*, (See Veliz-Cuba et al (2016) for the details of the derivation)

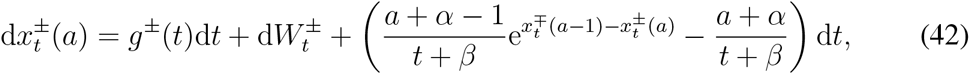

where 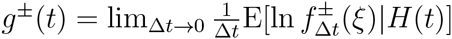 and *W^i^* satisfies 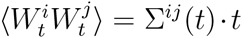 with 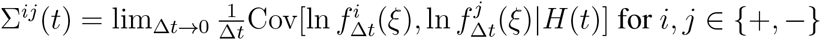.

Note that Eq. (42) is an infinite set of differential equations, one for each pair 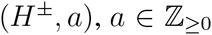. The initial conditions at *t* = 0 are given by *x*^±^(*a*) = ln P_0_(*H*^±^, *a*). To be consistent with the prior over the rate, *∊*, we can choose a Poisson prior over a with mean, *α*, i.e. 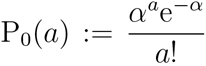. The initial conditions for Eq. (42) are given by 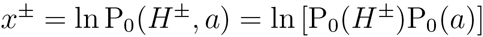. Note also that Eq. (42) at the boundary *a* = 0 is a special case. Since at those values there is no influx of probability from *a* – 1, Eq. (42) reduces to

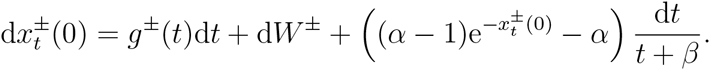

Lastly, note that we can obtain an evolution equations for the the likelihoods, 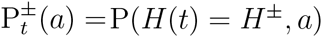, by applying the change of variables 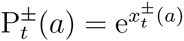. Itô’s change of coordinates rules (Gardiner, 2004) imply that Eq. (42) is equivalent to

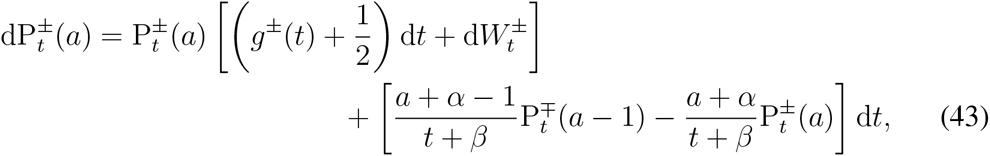

where now initial conditions at *t* = 0 are simply 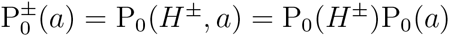. We will compare the full system, Eq. (43), with an approximation using a moment expansion in subsection 5.2.

#### Two states with asymmetric rates

Next we consider the case where the state of the environment, {*H_t_*}, is still a continuous-time Markov chain with state space {*H*^1^, *H*^2^}, but the probabilities of transition between the two states are asymmetric: P(*H_t+Δt_* = *H^i^*|*H_t_* = *H^j^*) = *∊^ij^*Δ*t* + *o*(Δ*t*), *i ≠ j*, where *∊*^12^ ≠ *∊*^21^. Thus, we must separately enumerate changepoints, 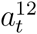 and 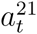, to obtain an estimate of the rates *∊*^12^ and *∊*^21^. In addition, we will rescale the enumeration of non-changepoints so that 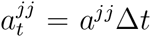, in anticipation of the divergence of *a^jj^* as Δ*t* − 0. This will mean the total *dwell time*, 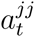, will be continuous, while the change point count will be discrete. The quantities 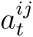 are then placed into a 2 × 2 matrix, 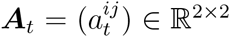, where 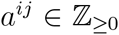 for *i* ≠ *j* and *a^jj^* ∈ ℝ*. Note that if the number of changepoints, 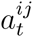, and the total dwell time in a state, 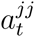, were known, the escape rate could be estimated as 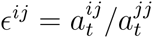.

As before, we will estimate the rate parameters, *∊^ij^*, using the posterior probability of the changepoint matrix, *a_t_*. We assume Gamma priors on each rate, so that *∊^ij^* ~ Gamma(*α_j_*,*β_j_*). By assumption the changepoint count, 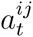, follows a Poisson distribution with parameter 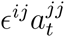, so that 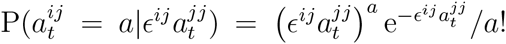.

Therefore, once 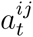 changepoints have been observed along with the dwell time 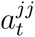, we have the posterior distribution 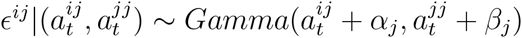, so

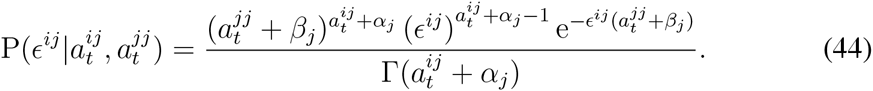

We now derive the continuum limit of Eq. (34). One key step of the derivation is the application of a change of variables to the changepoint matrix a, where we replace the non-changepoint counts with dwell times *t^j^*, defined as 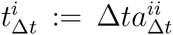 for Δ*t* = *t_n_* – *t_n−1_*. This is necessary, due to the divergence of 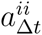 as Δ*t* → 0. In the limit Δ*t* → 0, the changepoint matrix becomes

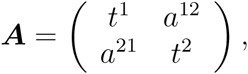

where *a^ij^* ∈ ℤ* is the changepoint counts from 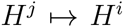, while *t_i_* ∈ ℝ* is the dwell time in state *H_i_*. Thus, taking logarithms, linearizing, and taking the limit Δ*t* → 0, we obtain the following system of stochastic partial differential equations (SPDEs) for the log likelihoods, 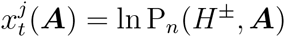:

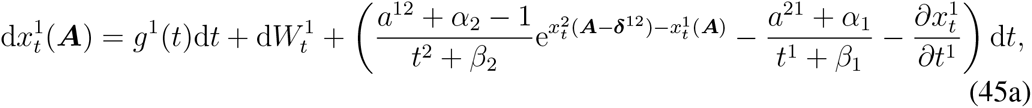

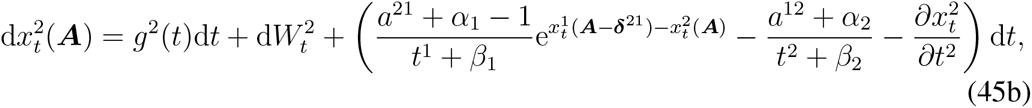

where the drift, *g^i^(t)*, and noise, *W^i^(t)*, are defined as before (for details, see Appendix 7.3). Note that the flux terms, 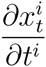, account for the flow of probability to longer dwell times *t^i^*. For example, the SPDE for 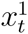 has a flux term for the linear increase of the dwell time *t*^1^, since this represents the environment remaining in state *H*^1^. These flux terms simply propagate the probability densities 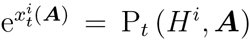 over the space (*t*^1^, *t*^2^), causing no net change in the probability of residing in either state 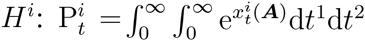.

Eq. (45) generalizes Eq. (34) as an infinite set of SPDEs, indexed by the discrete variables (*H^j^, a*^12^, *a*^21^) where *a*^12^, *a*^21^ ∈ ℤ_≥0_. Each SPDE is over the space (*t*^1^, *t*^2^), and it is always true that *t*^1^ + *t*^2^ = *t*. Initial conditions at *t* = 0 are given by *x^j^*(***A***) = ln[P_0_(*H^j^*) ° P_0_(***A***)]. For consistency with the prior on the rates, *∊^ij^*, we choose a Poisson prior over the changepoint counts *a^ij^, i* ≠ *j*, and a Dirac delta distribution prior over the dwell times *t^i^*

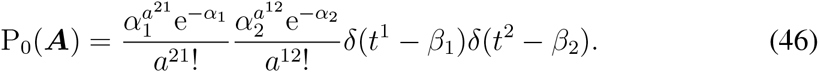

As before, Eq. (45) at the boundaries *a*^12^ = 0 and *a*^21^ = 0 is a special case, since there will be no influx of probability from *a*^12^ – 1 or *a*^21^ – 1.

As in the symmetric case, we can convert Eq. (45) to equations describing the evolution of the likelihoods 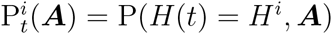. Applying the change of variables 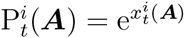, we find

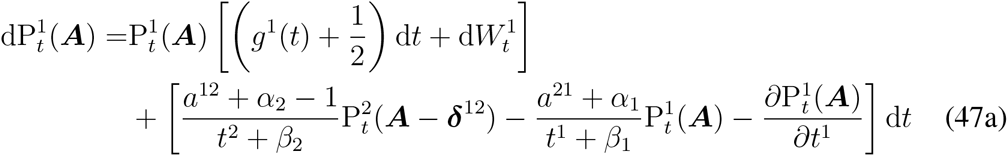

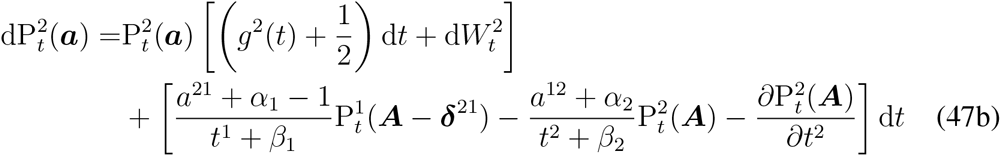

where now initial conditions at *t* = 0 are 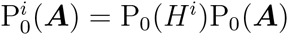.

#### Multiple states with symmetric rates

The continuum limit in the case of *N* states, {*H*^1^,…, *H^n^*}, with symmetric transition rates can be derived as with *N* = 2 (See Appendix 7.4 for details). Again, denote the transition probabilities by P(*H_t+Δt_* = *H^i^*|*H_t_* = *H^j^*) = *∊^ij^*Δ*t* + *o*(Δ*t*), and the rate of switching from one to any other state by *∊* = (*N* – 1)*∊^ij^*.

Assuming again a Gamma prior on the transition rate, *∊* ~ Gamma(*α,β*), and introducing 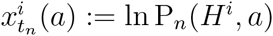, we obtain the SDE

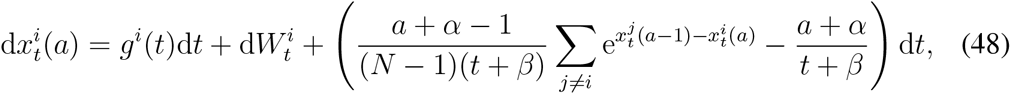

where 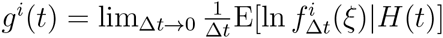 and *W^i^* satisfies 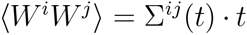 with 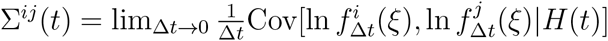.

Eq. (48) is again an infinite set of stochastic differential equations, one for each pair (*H^i^, a*), *i* ∈ 1,…, *N*, *a* ∈ ℤ_≥0_. We have some freedom in choosing initial conditions at *t* = 0. For example, since *x^i^(a)* = ln P_0_(*H^i^, a*), we can use the Poisson distribution discussed in the case of two states.

The posterior over the transition rate, *∊*, is

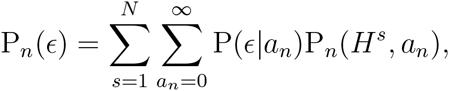

where P(*∊*|*a_n_*) is the Gamma distribution given by Eq. 35. Similar to Eq. 20, the expected rate is

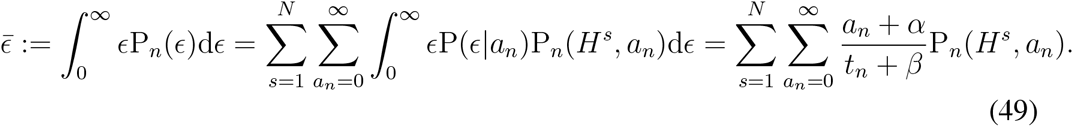

An equivalent argument can be used to obtain the posterior over the rates in the asymmetric case.

### 5.2 Moment hierarchy for the 2-state process

In the previous section, we approximated the evolution of the joint probabilities of environmental states and changepoint counts. The result, in the symmetric case, was an infinite set of SDEs, one for each combination of state and changepoint values (*H^i^, a*). However, an observer is mainly concerned with the current state of the environment. The changepoint count is important for this inference, but may not be of direct interest itself. We next derive simpler, approximate models that do not track the entire joint distribution over all changepoint counts, but only essential aspects of this distribution. We do so by deriving a hierarchy of iterative equations for the moments of the distribution of changepoint counts, *a* ∈ ℤ_≥0_, focusing specifically on the two state symmetric case.

Our goal in deriving moment equations is to have a low-dimensional, and reasonably tractable, system of SDEs. Similar to previous studies of sequential decision making algorithms (Bogacz et al, 2006), such low-dimensional systems can be used to inform neurophysiologically relevant population rate models of the evidence accumulation process. To begin, we consider the infinite system of SDEs given in the two state symmetric case, Eq. (43). Our reduction then proceeds by computing the SDEs associated with the lower order (0th, 1st, and 2nd) moments over the changepoint count a:

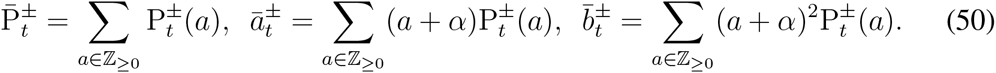

We denote the moments using bars 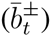. Below, when we discuss cumulants, we will represent them using hats 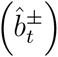. Note that the “0th” moments are the marginal probabilities of *H*^+^ and *H*^−^.

We begin by summing Eq. (43) over all *a* ∈ ℤ_≥0_ and applying Eq. (50) to find this generates an SDE for the evolution of the moments 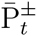 given

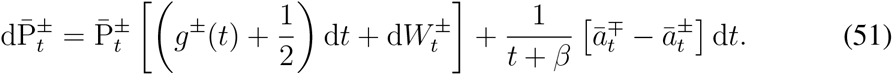

where we have used the fact that

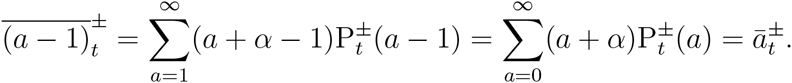

The SDE given by Eq. (51) for the zeroth moment, 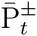, depends on the first moment, 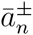. We can determine values for the first moment by either obtaining the next SDE in the moment hierarchy, or assuming a reasonable functional form for 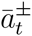. For instance, if the transition rate e is known we can assume 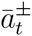, so that 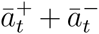 is approximately the mean of the counting process with rate *∊*. In this case, the continuum limit *t* − ∈ of Eq. (51) becomes

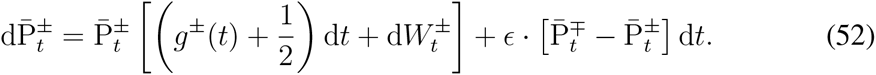

As expected, this is the two state version of Eq. (2), with known rate, *∊*. However, if the observer has no prior knowledge of the rate, *∊*, then 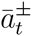 should evolve towards (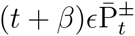 at a rate that depends on the noisiness of observations.

To obtain an equation for 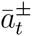 we multiply Eq. (43) by (*a* + *α*) and sum to yield,

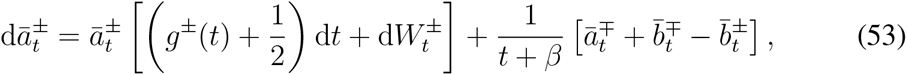

This equation relates the zeroth, first, and second moments, 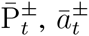, and 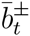. Again, we require an expression for the next moment, 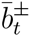, to close the system of Eqs. (51, 53). We could obtain an equation for 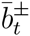 by multiplying Eq. (43) by (*a* + *α*)^2^ and summing. However, as is typical with moment hierarchies, we would not be able to close the system as equations for subsequent moments will depend on successively higher moments (Socha, 2007; Kuehn, 2016). To close the equations for 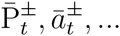 we can truncate: One possibility is cumulant-neglect (Whittle, 1957; Socha, 2007), which assumes all cumulants above a given order grow more slowly than the moment itself and can thus be ignored. This allows one to express the highest order moment as a function of the lower order moments, since a moment is an algebraic function of its associated cumulant and lower moments. For instance, neglecting the second cumulant 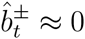 allows us to approximate the second moment as 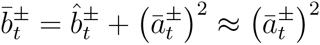.^3^

Applying cumulant-neglect to the second moment, 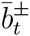, in Eqs. (51,53), using the change of variables, 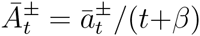, and the fact that 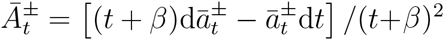, we obtain a closed system of equations for the zeroth and first moments,

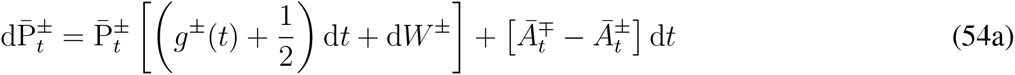

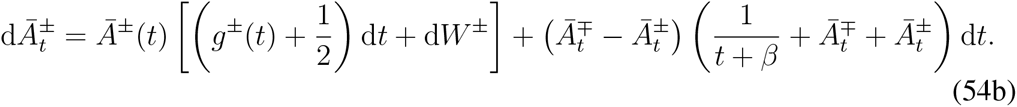

Here initial conditions are given by 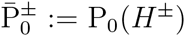 and 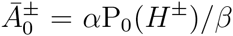. We show in Appendix 7.5 that Eq. (54) is also consistent with Eq. (2), which holds in the case of two states and known rate e. Trajectories of Eq. (54) are shown in Fig. 5. Note that both 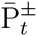 and 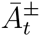 tend to increase when 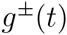 is the maximal drift rate, i.e. when *H*^±^ is the true environmental state. Thus, we expect that when 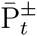 is high (close to unity) then 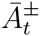 will tend to be larger than 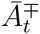.

**Figure 5:**
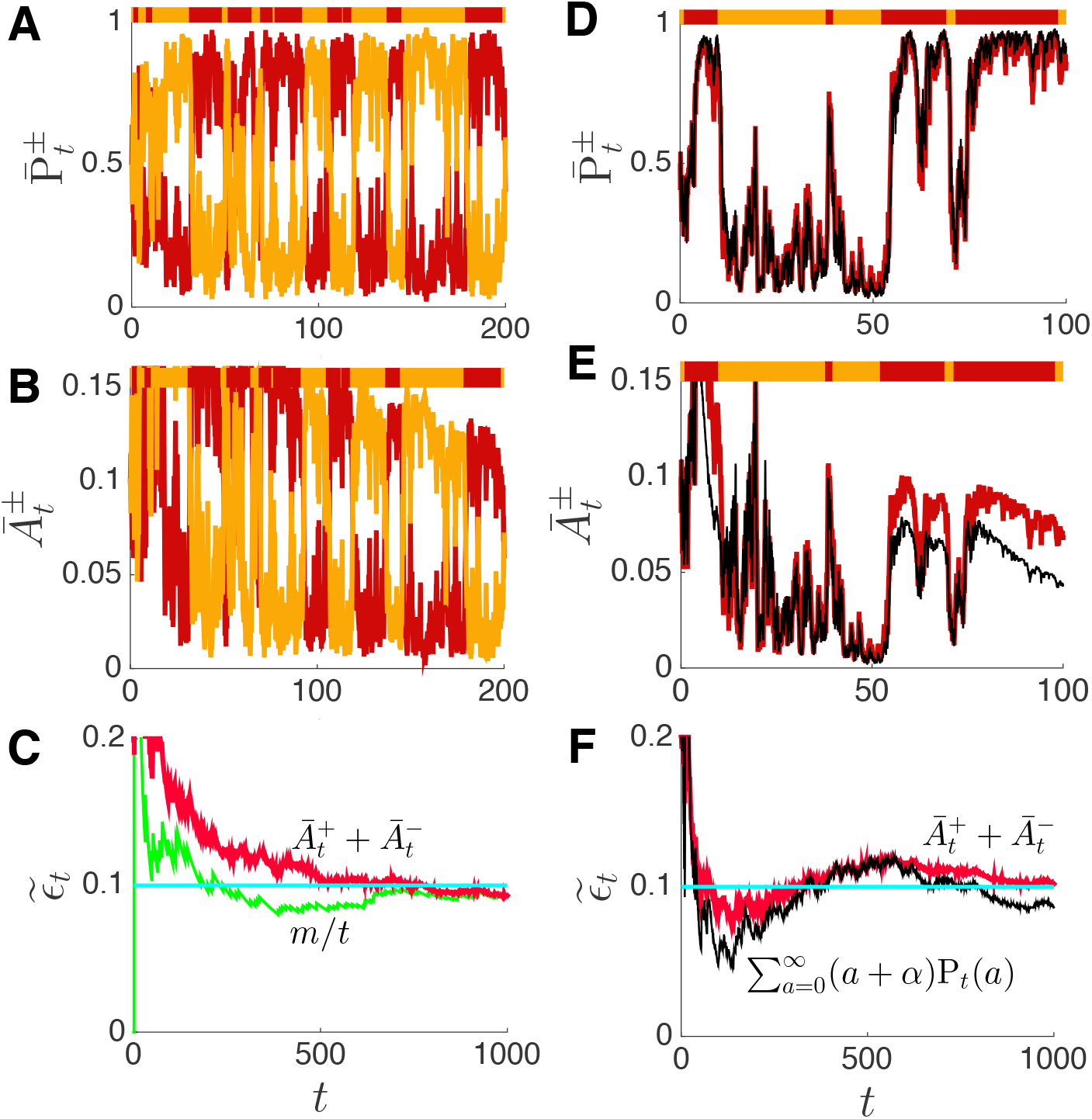
The dynamics of the first two moments, as approximated by Eq. (54). (A) The probabilities 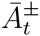 track the present state of the environment (bar above), switching with rate *∊* = 0.1, and approach the stationary densities around the equilibria determined by the dichotomous drift terms *g*^±^(*t*). (B) The first moments, 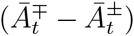, also switch with the environmental state and alternate between the neighborhoods of two points. (C) The sum 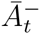 provides a running point estimate of the environmental transition rate, *∊*, as shown in Eq. (55). The estimate is determined by the actual changepoints, and noisily tracks *m/t*, where *m* is the actual number of changepoints. (D,E,F) Same as A,B,C, but the moment simulations are compared with numerical simulations of the full system of SDEs given by Eq. (43). (D) Thick red line is 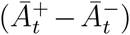 from Eq. (54) and thin black line is 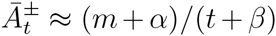 using Eq. (43); (E) Thick red line is 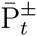 from Eq. (54) and thin black line is 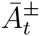 using Eq. (43); (F) Estimates of 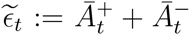 using Eq. (54) and Eq. (43). Details about the simulation method, initial conditions, and parameters are provided in Appendix 7.7.

Immediately after a changepoint (where the maximal drift rate *g*^±^(*t*) changes), there is an additional contribution to the increase of 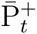 due to the 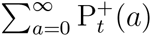 term. It is this brief burst of additional input to the subsequently dominant variable that generates the counting process, which enumerates changepoints. For instance, when a *H*^+^ ↦ *H*^−^ switch occurs, an increase in 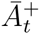 will temporarily be driven both by the drift term, *g*^−^(*t*), and the nonlinear term involving 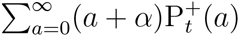. The burst of input generated by the nonlinear term in Eq. (54b) has an amplitude that decays nonautonomously with time. In fact, it can be shown that when the signal-to-noise ratio of the system is quite high, the variables 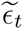, which is effectively the true changepoint count *m* divided by elapsed time as modified by the prior.

We can also obtain a point-estimate of the transition rate of the environment, which we define as 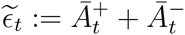, since the following relations hold:

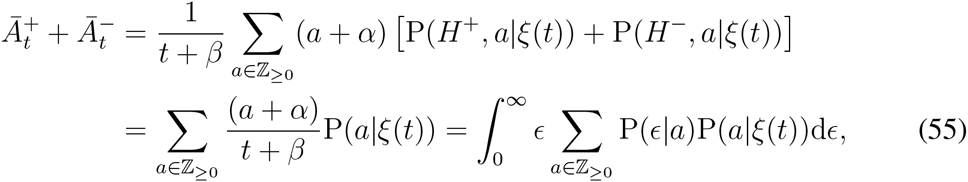

This estimate is an average over the distribution of possible changepoint counts, *a*, given the observations, *ξ(t)*. Here P(*∊*|*a*) is a Gamma distribution with parameters *α* and *β*. In Fig. 5C we compare this approximation, 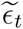, with the true change rate *∊* and the running estimate *m/t*, obtained from the actual number of changepoints, *m*.

In Fig. 5D,E these approximations are compared to Eq. (43), the full SDE giving the distribution over all changepoint counts, *a*. Notice that the first moments 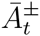 are overestimates of the true average, 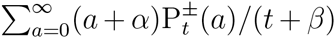, obtained from Eq. (43). We expect this is due to the fact that the moment equations, Eq. (54), tend to overcount the number of changepoints. Fluctuations lead to an increase in the number of events whereby 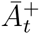 and 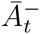 exchange dominance 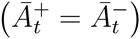, which will lead to a burst of input to one of the variables 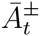. As a consequence, the transition rate tends to be overestimated by Eq. (54) compared to Eq. (43).

In sum, while the inference approximation given by Eq. (54) does not provide an estimate of the variance, it does provide insight into the computations needed to estimate the changepoint count and transition probability. Transitions increment the running estimate of the changepoint count, and this increment decays over time, inversely with the total observation time t. Similar equations for the moments can be obtained in the case of asymmetric transition rates, or more than two choices using Eq. (47) and Eq. (48) respectively, although we omit their derivation here.

## 6 Learning transition rate in neural populations with plasticity

Models of decision making often consist of mutually inhibitory neural populations with finely tuned synaptic weights (Machens et al, 2005; McMillen and Holmes, 2006; Wong et al, 2007). For instance, many models of evidence integration in two alternative choice tasks assume that synaptic connectivity is tuned so that the full system exhibits line attractor dynamics in the absence of inputs. Such networks integrate inputs perfectly, and maintain this integrated information in memory after the inputs are removed. However, in changing environments optimal evidence integration should be leaky, since older information becomes irrelevant for the present decision (Deneve, 2008; Glaze et al, 2015).

We previously showed that optimal integration in changing environments can be implemented by mutually excitatory neural populations (Veliz-Cuba et al, 2016). Instead of a line attractor, the resulting dynamical systems contain globally attracting fixed points. Such leaky integrators maintain a limited memory of their inputs on a timescale determined by the frequency of environmental changes. However, in this previous work we assumed that the rates of the environmental changes were known to the observer. Here, we show that when these rates are not known a priori, a plastic neu-ronal network is capable of learning and implicitly representing them through coupling strengths between neural populations.

### Symmetric environment

We begin with Eq. (54), the leading order equations for the likelihood 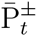 and change rate variables 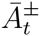 derived in Section 5.2. We interpret the likelihoods as neural population activity variables 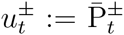, reflecting a common modeling assumption that two populations receive separate streams of input associated with evidence for either choice (Bogacz et al, 2006). Next, we define a new variable 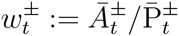, which represents the synaptic weight between these neural populations. In particular, 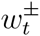 represents the strength of coupling from 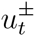 to 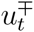 as well as the local inhibitory coupling within 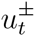. Applying this change of variables to Eq. (54), we derive a set of equations for the population activities, 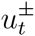, and their associated synaptic weights, 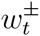:

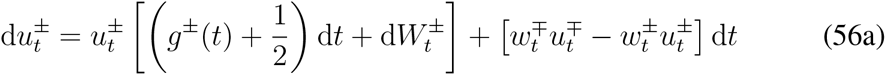

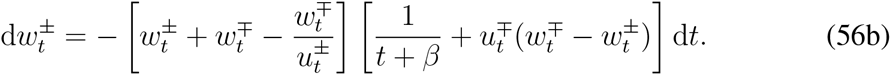

This is a neural population model with a rate-correlation based plasticity rule (Miller, 1994; Pfister and Gerstner, 2006). We can interpret the non-autonomous term, 1 /(*t*+*β*), as modeling the dynamics of a chemical agent involved in the plasticity process whose availability decays over time. Simple chemical degradation kinetics for a concentration *C_t_* yield such a function when

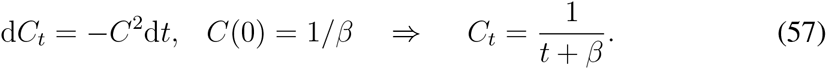

We briefly analyze the model, Eq. (56), by considering the limit of no observation-noise. That is, we assume *g*^±^(*t*) → ±∞ when *H_t_* = *H*^+^ and Σ^++^ → 0, where 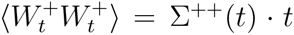, and analogous relations hold when *H_t_* = *H*^−^. As a result, when *H_t_* = *H^+^*, then 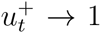 and 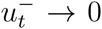, which we demonstrate in Appendix 7.6. Plugging the expressions 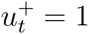 and 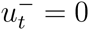 into Eq. (56b) for 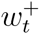, we find

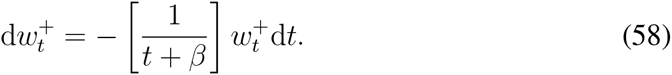

Next, we write Eq. (56b) for 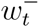 in the form

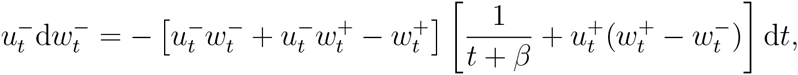

so by plugging in 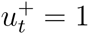 and 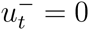, we find 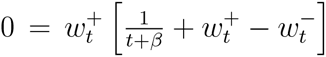, which, when 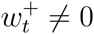, simplifies to

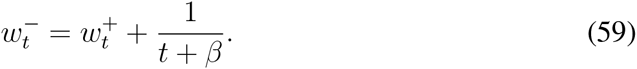

An analogous pair of equations holds when *H_t_* = *H^−^* and thus 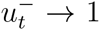 and 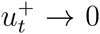. Solving Eq. (58,59) and their *H_t_* = *H^−^* counterparts iteratively, we find that in the limit of no observation-noise (e.g., *g*^±^(*t*) → ±∞ when *H_t_* = *H^±^*),

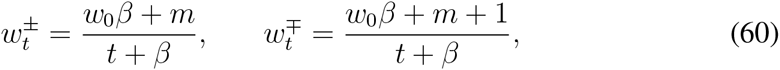

where *H_t_* = *H^±^* and *w*_0_ := *w^j^* (0) for *H*(0) = *H^j^*, so *w*_0_ constitutes the initial estimate of the change rate of the environment. Here, *m* is the number of changepoints in the time series *H_t_* during the time interval [0, *t*]. Eq. (60) can be re-expressed in the form of a rate-based plasticity rule

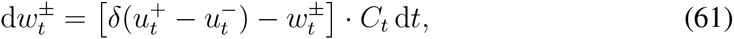

where *δ*(*u*) is the Dirac delta distribution, along with Eq. (57) for the concentration decay of the agent *C_t_*. Note, that the non-negative term 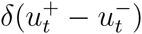 in Eq. (61) results in long term potentiation (LTP) of both synaptic weights, 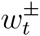, whenever the neural activities, 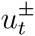, are both high, *i.e*. when their values cross at 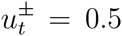. During such changes, the weights 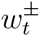 are incremented. Outside of these transient switching epochs, there is a constant long term depression (LTD) of the synaptic weights 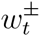 modeled by the term 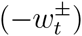.

We schematize (Fig. 6A) and simulate (Fig. 6B,C) the resulting plastic neural population network:

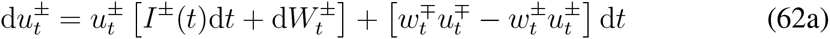

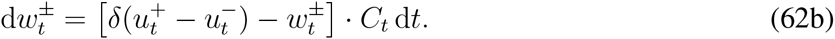

**Figure 6:**
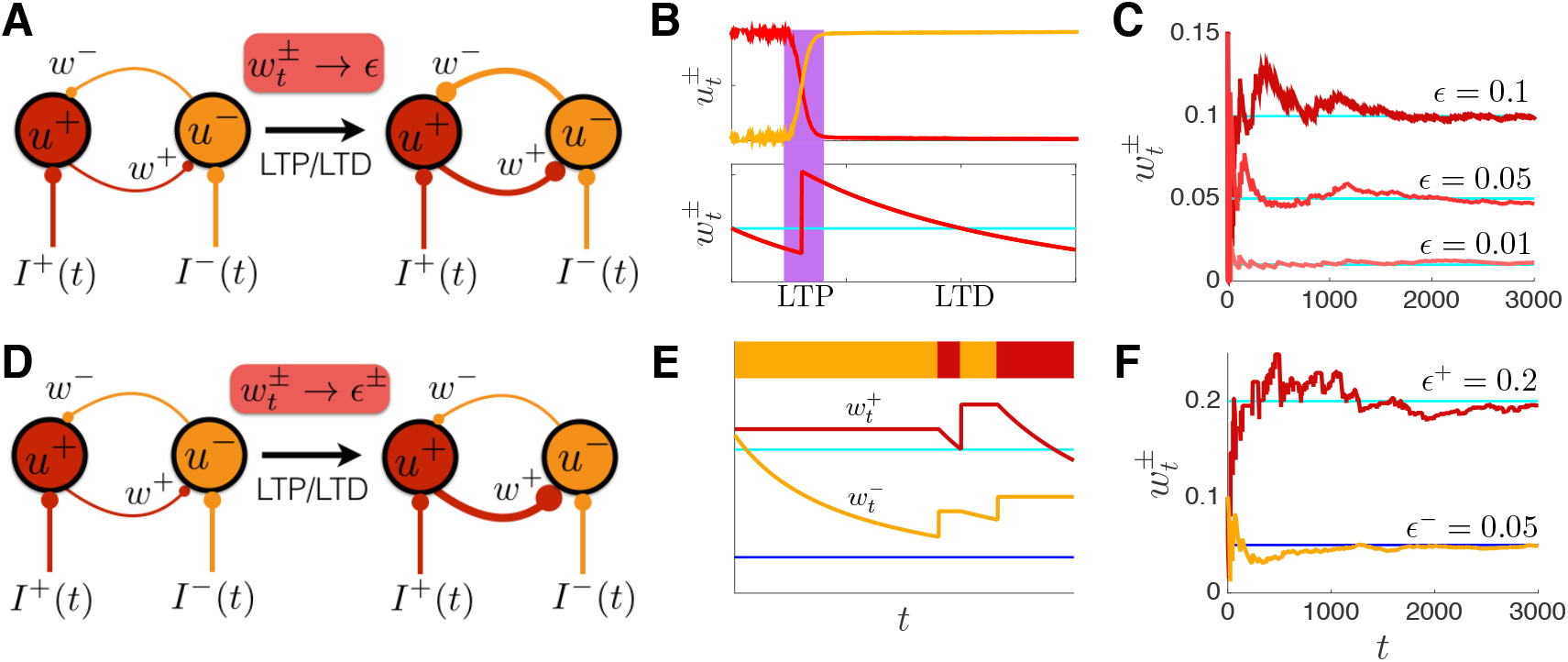
Neural network model with plasticity, inferring the current state Ht and rates *∊*^±^ of environmental change. (A) Schematic showing the synaptic weight *w*^±^ from neural population *u*^±^ ↦ *u*^∓^ evolving through long term potentiation (LTP) and long term depression (LTD) to match the environment’s rate of change, *∊*^±^ := *∊*. (B) When the neural populations exchange dominance, their activity levels *u*^±^ are both transiently high. As a result, both synaptic weights, *w*^±^, increase via LTP. When only one population is active, both weights decay via LTD, as described by Eq. (62b). (C) Inference of the rate, *∊*, via long term plasticity of the weights for *∊* = 0.01, 0.05, 0.1. Though the signal-to-noise ratio is finite (See Appendix 7.7), the weights in the network described by Eq. (62) converge to the actual rate, *∊*. (D) Schematic showing the evolution of weights 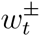 when rates are asymmetric, *∊*^+^ > *∊*^−^, so that 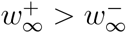,. The network is described by Eq. (65). (E) Only the weight 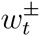 decays through LTD when population 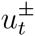 is active, and only the weight 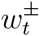 is potentiated through LTP when dominance switches from 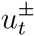 to 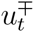, as in Eq. (65). (F) Network weights 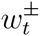 converge to the asymmetric rates, *∊*^±^. See Appendix 7.7 for details about the simulations.

The constant 1/2 has been absorbed into the population input, so that *I*^±^ (*t*) = *g*^±^(*t*) + 1/2. It is important to note that Eq. (62) only performs optimal inference in the limit of no observation-noise. Perturbing away from this case, we expect the performance to be sub-optimal. However, as can be seen in Fig. 6C, the correct change rate is approximated reasonably well.

Eq. (62), thus models evidence accumulation in a symmetrically changing environment when the change rate, *∊*, is not known a priori. The model is based on the recursive equation for the joint probability of the environmental state, *H*^±^, and changepoint count, a, derived in Section 3.1. We obtained a tractable model by first passing to the continuum limit, and then applying a moment closure approximation to reduce the dimension of the resulting equations. Obtaining the low-dimensional approximation in Eq. (54) was crucial to obtaining a neural population model that approximates state inference. We next extend this model to the case of asymmetric rates of change.

### Asymmetric environment

The continuum limit of the inference process in an asymmetric environment, Eq. (47), provides several pieces of information we can use to identify an approximate neural population model. First, under the assumption of large signal-to-noise ratios, the synaptic weights should evolve to reflect the number of detected change-points, rescaled by the amount of time spent in each state

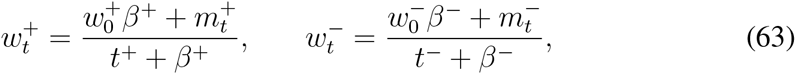

where 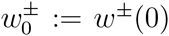 are the network’s initial estimates of the change rates *∊*^±^, *m*^±^ is the true number of changepoints *H*^±^ ↦ *H*^∓^ during the time interval [0, *t*], and *t*^±^ is the total length of time spent in the state *H*^±^.^4^ Second, the flux term in Eq. (47) indicates that a memory process is needed to store the estimated time *t*^±^ spent in each state *H*^±^. This can be accomplished by modifying Eq. (57) for the plasticity agent, so that each decays only when the neural population of origin, 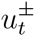, is active. Thus we obtain the pair of equations:

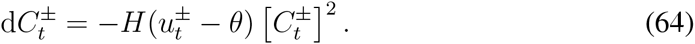

Expressing Eq. (63) as a system of equations for the synaptic weights, 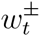, yields:

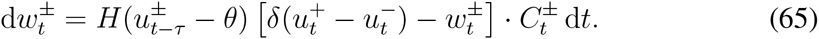

Here the function *H*(*u*^±^(*t* – *τ*) – *θ*) for *θ* ≥ 0.5, and *τ* > 0 enforces the requirement that the population 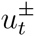 must have a high rate of activity prior to the LTP event. Thus, to learn asymmetric weights, there should be a small delay *τ* accounting for the time it takes for the presynaptic firing rate to trigger the plasticity process (Gütig et al, 2003). We demonstrate the performance of the network whose weights evolve according to Eq. (65) in Fig. 6D,E. Note, the network with weights evolving according to Eq. (65) can still infer symmetric transition rates *∊*^±^ = *∊*, but it will do so at half the rate of the network Eq. (62). This is due to the fact that Eq. (65) counts changepoints and dwell times of each state *H*^±^ separately.

We have thus shown that the recursive update equations for the state probability in a dynamic environment lead to plausible neural network models that approximate the same inference. Previous neural network models of decision making have tended to interpret population rates as a representation of posterior probability (Bogacz et al, 2006; Beck et al, 2008). We have shown that the synaptic weight between populations can represent the change rate of the environment. As a result, standard rate-correlation models of plasticity can be used to implement the change rate inference process.

## 7 Discussion

Evidence integration models have a long history in neuroscience (Ratcliff and McKoon, 2008). These normative models conform with behavioral observations across species (Brunton et al, 2013), and have been used to explain the neural activity that underpins decisions (Gold and Shadlen, 2007). However, animals make decisions in an environment that is seldom static (Portugues and Engert, 2009). The relevance of available information, the accessibility, and the payoff of different choices can all fluctuate. It is thus important to extend evidence accumulation models to such cases.

We have shown how ideal observers accumulate evidence to make decisions when there are multiple, discrete choices, and the correct choice changes in time. We assumed that the rates of transition between environmental states are initially unknown to the observer. An ideal observer must therefore integrate information from measurements toconcurrently estimate both the transition rates and the current state of the environment. Importantly, these two inference processes are coupled: Knowledge of the rate allows the observer to appropriately discount older information to infer the present state, while knowledge of transitions between states is in turn necessary to infer the rate.

Inference when all transition rates are identical is straightforward to implement in resulting models. An ideal observer only needs to track the probability of the environmental state and the total changepoint count, regardless of the states between which the change occurred. However, when the transition rates are asymmetric, the resulting models are more complex. In this case, an ideal observer must estimate a matrix of changepoint counts, distinguished by the starting and ending states. The number of possible matrices grows polynomially with the number of observations. This computation is difficult to implement, and we do not suggest that animals make inferences about environmental variability in this way. However, understanding the ideal inference process allowed us to identify its most important features. In turn, we derived tractable approximations and plausible neural implementations, whose performance compared well with that of an ideal observer (Fig. 5D,E,F). We believe humans and other animals do generally implement approximate strategies when they need to infer such rates (Lange and Dukas, 2009). Ideal observer models allow us to understand what inferences can be made with the available information, which assumptions of the observer are important (e.g., assuming an incorrect transition rate does not always have a large impact on performance), and how such inferences could be approximated in networks of the brain and other biological computers.

In many naturally occurring decisions like foraging, mate selection, and home-site choice, animals simply need to identify the best alternative rather than the rate of environmental change (Johnson et al, 2013). Therefore, rapid approximations, or a guess of the environmental change rate may provide better initial performance than learning the rate, which could be slow. Moreover, it appears that when measurements are noisy, rates cannot be learned precisely even in the limit of infinite observations. Thus, learning the rate may only be useful when noise is too high for single measurements to determine the correct alternative, but sufficiently low to make rate inference possible. There is evidence that humans adjust their rate of evidence-discounting, based on the actual changerate of the environment (Glaze et al, 2015). However, further psychophysical studies are needed to identify whether subjects use heuristic strategies to learn or something close to the normative models we derived here.

A number of related models have been developed previously (Wilson et al, 2010; Adams and MacKay, 2007). The present model is somewhat different, as a finite number of choices implies that the present environmental state is dependent on the previous state. As a result, we found it was more efficient to implement an update equation that estimated the present environmental state and the changepoints, rather than the time in the present state.

Several of the assumptions we have made in this study could be modified to extend our analysis to more general situations. For instance, we have assumed that the observer’s eventual choice does not affect the environment. However, in many natural situations changes in the environment are a consequence of the observer’s actions (Cisek and Pastor-Bernier, 2014). In more realistic situations it is likely that there is a sequence of actions leading to an ultimate decision, and each action can influence the information available to the observer. An animal making a foraging decision in a group collects more evidence once it moves toward a particular food patch, but it may also draw other members with it, changing the subsequent availability of food there (Petit et al, 2009). Thus including a sequence of actions, and their impact on the available information and the environment would be necessary in a realistic model. Another possibility is that changes to the environment are non-Markovian and/or involve multiple timescales. Extending our ideal observer models to estimate such change statistics might require derivation of multi-step update equations. In such cases, we expect the truncations we have applied in this work would be useful for identifying tractable approximations of the optimal inference process.

Optimal models of evidence accumulation are useful both as baselines to compare to performance in psychophysical experiments, and starting points for identifying plausible neuronal network implementations. Our core contribution here has been to present a general model of evidence accumulation in a dynamic environment, when an observer has no prior knowledge of the rate of change. An unavoidable feature of these models is that the number of variables the observer must track grows as more observations are made, and growth is more rapid in asymmetric environments with multiple environmental states. This motivated our development of continuum approximations and lowdimensional moment equations for the optimal models, which suggest more plausible neural computations. We hope this work will foster future theoretical studies that will extend this framework, as well as experiments that could validate the models herein. To fully understand the neural mechanisms of evidence accumulation, we must account for the wide variety of conditions that organisms encounter when making decisions.

## Acknowledgments

Funding was provided by NSF-DMS-1517629 (AER, KJ, andZPK); NSF-DMS-1311755 (ZPK); and NSF/NIGMS-R01GM104974 (KJ).

## Appendix

### 7.1 Two-state system with unknown symmetric rate

We show how to derive Eq. (3) from the main text. Bayes’ rule and the law of total probability first yield:

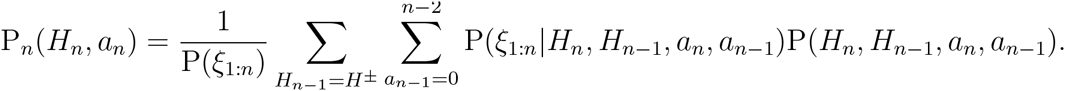

Using the conditional independence of observations,

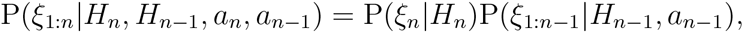

we find that,

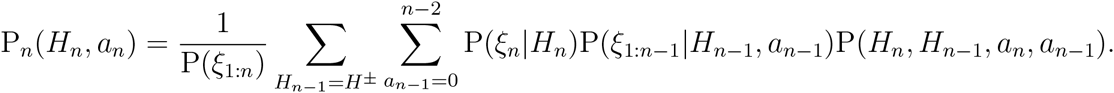

Furthermore, we can use the definition of conditional probability to write,

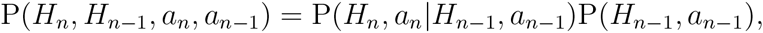

and Bayes’ rule also implies,

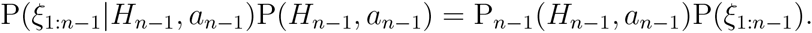

Hence, we derive Eq. (3) from the main text,

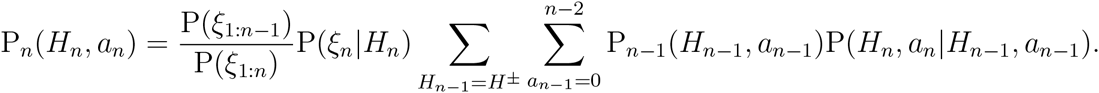

### 7.2 Numerical methods for free response protocol

The free response protocol is simulated by evolving the update Eq. (17) and subsequently computing the log likelihood ratio *L_n_* := log(*R_n_*) using Eq. (18) at each timestep *n*. Each point along the curves in Fig. 3C corresponds to an average waiting time and average performance corresponding to a threshold value *θ* over 100,000 simulations. For each value of *θ*, the simulation is terminated when |*L_n_*| > *θ* and the choice is given by the sign of *L_n_*. To avoid excessively long simulations, we removed any that lasted longer than *n* = 5000, but we found changing this upper bound did not affect averages considerably. There were 400 values of *θ* chosen, discretizing the interval from *θ* = 0 to *θ* = 3.89.

### 7.3 Continuum limit for two states with asymmetric rates

We begin by considering Eq. (34a), which provides an update of the probability of being in state *H*^1^ after *n* observations, given the specific changepoint matrix *α*:

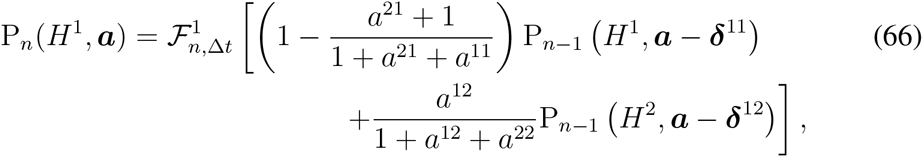

where we have defined 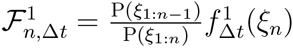. Subsequently, we divide by P_*n*−1_(*H*^1^, *α*) and take the logarithm to find:

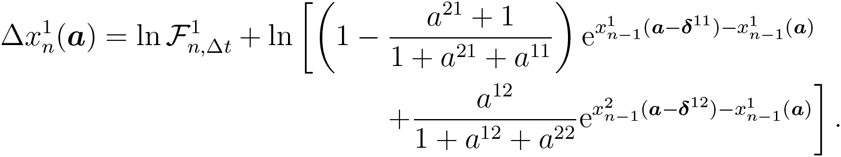

Now, as we are taking the continuum limit, we consider 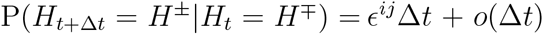, where 0 ≥ *∊^ij^* < ∞. Given a *ℝ*^2x2^ transition rate matrix with off-diagonal entries *∊^ij^*Δ*t* + *o*(Δ*t*) and diagonal entries 1 – *∊^ij^*Δ*t* + *o*(Δ*t*), the expected changepoint and non-changepoint counts after *n* timesteps will be 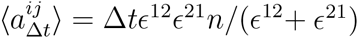 and 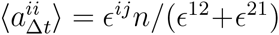. Thus, while the changepoint counts 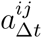 scale with Δ*t*, the non-changepoint counts do not. In the continuum limit, we will choose a time *t* := *n*Δ*t* and take Δ*t* → 0 while keeping *t* constant, so the number of timesteps diverges like *n* = *t*/(Δ*t*) for a fixed time *t*. While the expected changepoint counts 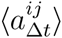 thus remain fixed, the non-changepoint counts 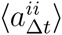 will diverge as (Δ*t*)^−1^, suggesting we should rescale non-changepoint counts to the absolute dwell time 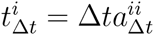. The expected value of the dwell times is then finite in this limit 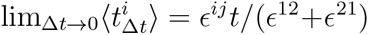. Performing this change of variables, we then define the changepoint matrix as involving changepoint counts *a^ij^* on the off-diagonal and dwell times along the diagonal: 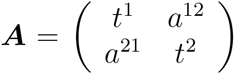 so an increment of non-changepoint count *a^ii^* now takes the form ***A*** + Δ*tδ^ii^*. As such, we can now expand:

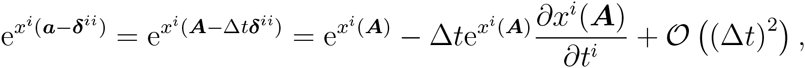

via application of the chain rule and noting *dt^i^*/*da^ii^* = Δ*t*. Note, we cannot perform such an expansion in Δ*t* to *x^i^*(***A*** – ***δ***^*ij*^), since perturbations to the matrix ***A*** in this case are 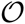(1). Truncating Eq. (66) to terms of 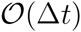 and incorporating the Poisson-delta prior Eq. (46), we find the discrete update equation becomes:

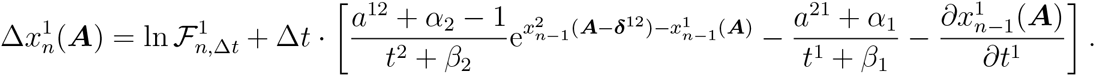

Lastly, upon taking the continuum limit Δ*t* → 0, we find that

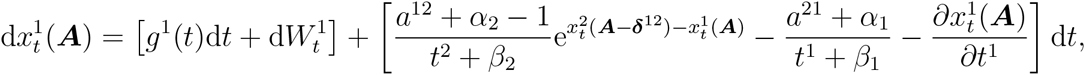

where the statistics of the drift *g*^1^(*t*) and noise 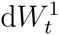 are analogous to those given after Eq. (42), only the transition rates of *H_t_* from *H^j^* ↦ *H^i^* are now *∊*^*ij*^. Note, due to the flux term 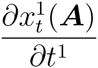 and continuum values for *t^j^ ∊* ℝ*, this is a stochastic partial differential equation (SPDE). An analogous SPDE can be derived for 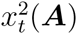 in the same way.

### 7.4 Continuum limit with multiple states and symmetric rates

The derivation parallels that with two states. To obtain the continuum limit, we use the generalized version of Eq. (3),

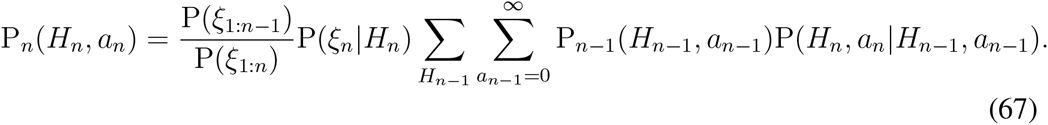

Assuming again a Gamma prior on the transition rate, *∊* ~ Gamma(*α, β*), and following the derivations of Eq. (23) and (38), we obtain

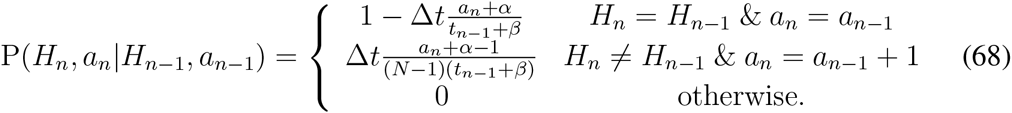

Using Eq. (68) in Eq. (67) yields

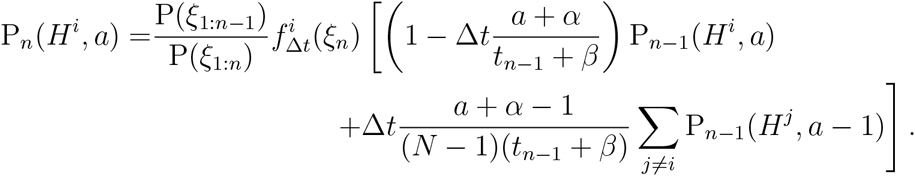

Dividing by *P*_*n*−1_(*H^i^,a*), taking logarithms, and denoting 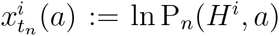 we obtain

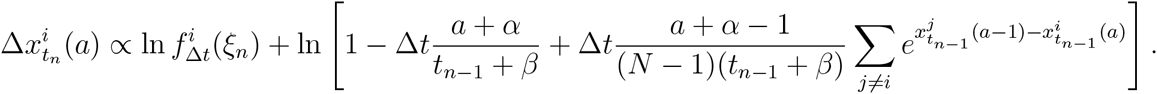

Using the approximation ln(1 + *z*) ≈ *z* valid for small *z* yields

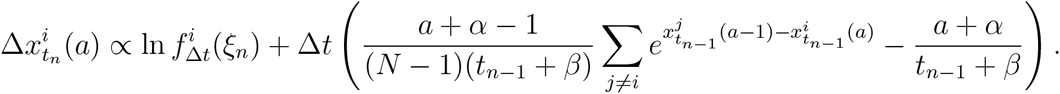

Similar to the *N* = 2 case, we may then take the continuum limit to yield Eq. (48).

### 7.5 Consistency of the moment hierarchy equations

We begin by taking the SDE given by Eq. (2) for *N* =2 states, and the known rate e, and changing variables to 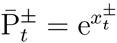, so

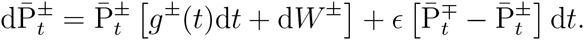

Furthermore, note that in the limit *t* → ∞, the 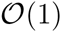 terms and 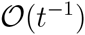 terms vanish in Eq. (54):

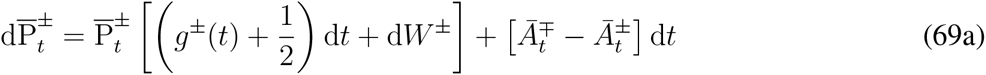

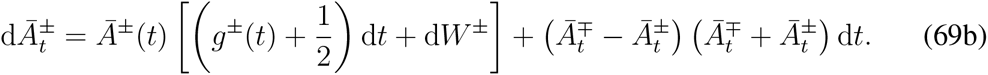

Therefore, in the event that 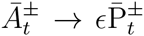 in the long time limit (*t* → ∞), we find the truncated system, Eq. (69), becomes

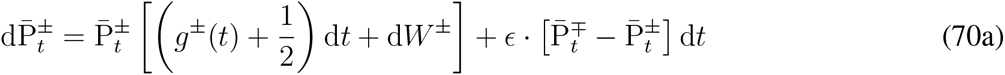

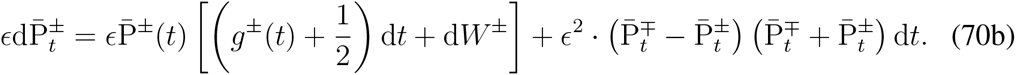

Dividing by *∊* and noting that 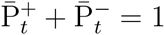, Eq. (70b) becomes

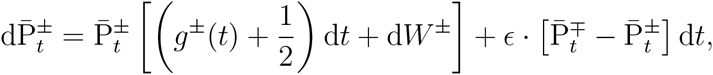

which is consistent with Eq. (70a), and indicates the truncated moment hierarchy is consistent with the case of known rates and two choices (*N* = 2) in the SDE Eq. (2).

### 7.6 Noise-free limit of the neural population model

Consider the neural population Eq. (56a) for the evolution of 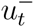 in the event of environmental state *H_t_* = *H*^+^ and no observation-noise 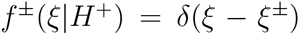. As a result, the drift terms diverge 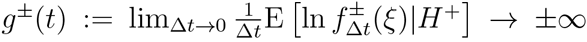 and the covariance matrix 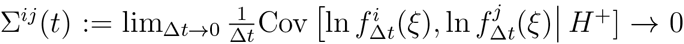. Thus, the dominant terms on the right hand side of Eq. (56a) for 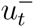 come from the input so 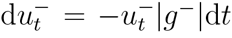, and the population activity immediately decays to 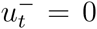. As a result, since 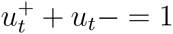, we expect 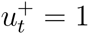, when *H_t_* = *H*^+^.

### 7.7 Numerical simulations of SDE models

Stochastic differential equation (SDE) models of evidence accumulation in symmetric environments changing between two states are simulated using a standard Euler-Maruyama integration algorithm (Higham, 2001). Eq. (43) describes the evolution of an infinite number of SDEs over the changepoint vector *a ∊* ℤ_≥0_, state vector *H_t_ ∊* {*H*^+^, *H*^−^}, and time *t* ∈ [0, *T*], so we truncate this space to *a* ∈ {0,1, 2,…., 1000}, which is sufficient for transition rates e and total simulation times *T* not too large. We compared our results to cases with longer state vectors *a* ∈ {0,1, 2,…., *a*_max_} and the changes were negligible. Simulations shown in Fig. 5 had a transition rate of *∊* = 0.1 and total run time of *T* = 1000 with timestep dt = 10^−3^. Observations were sampled from a normal distribution 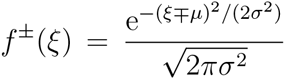 with mean *μ* = 0.5 and variance *σ*^2^ = 1 so the signal-to-noise ratio was 2*μ*/*σ* = 1. Initial conditions were chosen so that 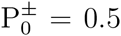 and 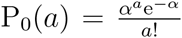 where *α* =1 and *β* = 5. A similar approach was used to numerically simulate the neural population model Eq. (62) and its variants to produce Fig. 6, but *μ* =1 and *σ* = 0.1, so the signal-to-noise ratio was 20.

## Footnotes

1 Equal spacing Δ*t* = *t_j_* − *t*_*j*−1_ is not necessary for all *j* = 2, …, *n*, but it does allow for a more concise derivation of the continuum limit. Irregular spacings would require a more careful selection of the scaling of the log likelihoods ln *f*^^±^^(*ξ*).

2 Note, we drop the ln [P(*ξ*_1:*n−i*_)/P(*ξ*_1:*n*_)] term below since it is common to all evolution equations. For determining the most likely option, only the relative magnitudes of the log likelihoods are important. In numerical simulations, we normalize to account for this discrepancy.

3 Note, we have used a hat to distinguish cumulants 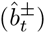, whereas bars still denote moments 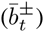.

4 We use the notation *H*^±^ for the two states here, for convenience and consistency with Eq. (62). Similarly, we use *∊*^±^ and *t*^±^ rather than the numerical notation of the asymmetric case in Section 5.1.

